# PRMT5 is Frequently Upregulated and a Potential Therapeutic Target in MTAP-deficient Malignant Peripheral Nerve Sheath Tumors

**DOI:** 10.64898/2026.03.09.710638

**Authors:** Dingxun Wang, Melissa L. Fishel, Azadeh Samiei, Silpa Gampala, Changdeng Hu, Shaoxiong Chen, GuangJun Zhang

## Abstract

Malignant peripheral nerve sheath tumors (MPNSTs) are highly aggressive sarcomas with poor prognosis and a strong tendency for metastasis and relapse. Surgical removal remains the mainstay of treatment but is frequently ineffective or impractical. Currently, no effective targeted therapy exists for this type of malignancy. PRMT5 has recently emerged as a promising therapeutic target in various human cancers with MTAP loss, which results in cancer cell dependency on PRMT5 activity. The frequent loss of MTAP in MPNSTs suggests that PRMT5 inhibition is a promising therapeutic option and enables the stratification of cancer patients with few treatment options. We first examined human nerve sheath tumor samples and found that increased PRMT5 expression and activity correlated with MTAP loss in 86.8% (33/38) of MPNSTs and in atypical neurofibromatous neoplasm with uncertain biologic potential (ANNUBP) (5/5). When PRMT5 activity was inhibited genetically and chemically, the cell growth of MTAP-deficient MPNST cell lines was suppressed, but not that of MTAP-proficient MPNST cell lines. Moreover, in the PRMT5-inhibited MTAP-deficient MPNST cell lines, spontaneous DNA damage accumulation was observed following G2/M cell cycle arrest. The DNA replication stress marker RPA32 decreased, and CHK1 was activated early after PRMT5 knockdown, likely contributing to the accumulation of DNA damage. In addition, we combined PRMT5 inhibition with the DNA-damaging agents doxorubicin and gemcitabine, resulting in synergistic effects and increased cancer cell death in MTAP-deficient MPNST cell lines. Together, these findings identify PRMT5 as a compelling therapeutic target in MTAP-deficient MPNSTs. This PRMT5 inhibition strategy has strong translational potential for MPNSTs.

**GRAPHICAL ABSTRACT:** 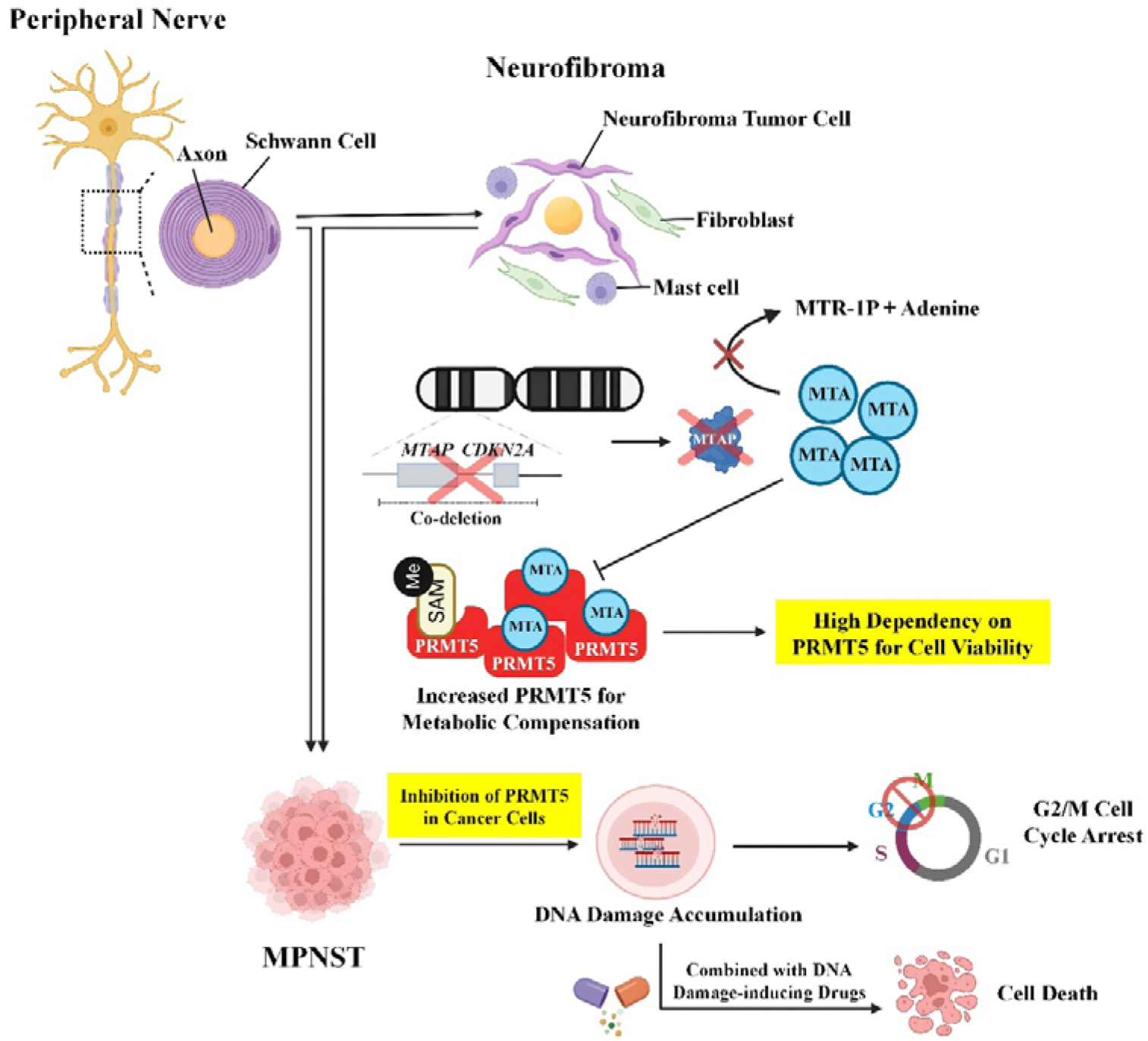

## INTRODUCTION

Malignant peripheral nerve sheath tumors (MPNSTs), also known as neurofibrosarcoma or neurosarcoma, originate from Schwann cells or the neural crest pluripotent cells ^1,2^. MPNSTs are highly aggressive tumors, accounting for 3-10% of soft tissue sarcomas ^3^. Etiologically, approximately half of patients have neurofibromatosis type 1 (NF1), and the other half are sporadic, likely due to different gene mutations, including TP53, NF2, and PRC2 ^4,5^. Despite MPNST being genomically heterogeneous, a few genetic events have been identified as critical for the malignant transformation of MPNST, whether arising from NF1 or *de novo* tumorigenesis. Homozygous loss of *CDKN2A* was most frequently observed in MPNST cases (69-80%), a finding that, notably, implicates it in the stepwise progression of MPNST ^6,7^. The *CDKN2A* gene, located on chromosome 9p21, encodes two critical tumor suppressor proteins, p14/ARF and p16/INK4A, which activate downstream tumor suppressors TP53 and RB1, respectively ^8,9^. The approximate 5-year survival of MPNST patients is around 40% due to their invasive nature and limited treatment options ^10,11^. MPNSTs are generally resistant to chemotherapy and radiotherapy ^2,12–14^, with surgical resection as the mainstay of treatment; however, this is often compromised by factors such as distant metastasis, local damage, and inoperable locations ^13^. Currently, no effective targeted therapies are available for this type of malignancy, and standard of care regimens such as doxorubicin plus ifosfamide are often unsuccessful ^15^. Thus, effective treatment strategies are highly desired.

Targeted cancer therapies are usually developed based on oncogene gain-of-function mutations, such as BRAF and NRAS, using chemical inhibitors or antibodies ^16^. In contrast, tumor suppressor genes are more complex to target. However, they also play equally important roles and are commonly deleted across many types of cancer. One example of the successful application of exploiting tumor suppressor gene expression is the use of PARP inhibitors for BRCA-mutant ovarian cancers, demonstrating the principle of synthetic lethality ^17,18^. Collateral lethality is a special case of synthetic lethality, in which an essential metabolite gene is co-deleted with a neighboring tumor suppressor gene ^17,19,20^. CDKN2A-MTAP-PRMT5 is an example of an exploitable collateral lethality in cancer ^21–23^. There is a high rate of homozygous deletion of *CDKN2A* in many types of cancer, including MPNST. The *CDKN2A* deletion often hijacks *MTAP*, a neighboring gene that encodes methylthioadenosine phosphorylase ^22,23^. MTAP participates in the salvage pathway of adenine/methionine. MTAP catalyzes the phosphorolysis of the substrate 5’-deoxy-5’-methylthioadenosine (MTA), a by-product of polyamine synthesis from S-adenosyl methionine (SAM), and produces adenine and methylthioribose-1-phosphate (MTR-1-P) for ATP and methionine recycling ^24,25^. The loss of MTAP impairs cell metabolism, leading to MTA accumulation and rendering cancer cells highly dependent on PRMT5 activity. Thus, inhibiting PRMT5 specifically blocks MTAP-deficient (low expression, loss, or loss of function) cancer cell growth ^22,23^. To be consistent and convenient, we use the terms *MTAP*-deficient and MTAP-proficient throughout this manuscript.

PRMT5 (protein arginine methyltransferase 5) is the major SAM-utilizing methyltransferase that catalyzes symmetric di-methylation on many substrates, including histones, cell signaling factors, transcriptional regulators, and mRNA splicing factors ^26,27^. Thus, PRMT5 is critical for several cellular activities. Recently, multiple studies have reported the pivotal role of PRMT5, with significantly elevated expression levels, in tumorigenesis and in the progression of many human cancers. Consistently, increased PRMT5 expression and activity have been widely reported to correlate with poorer prognostic outcomes ^27,28^.

Frequent deletions of *CDKN2A* (∼80%) and *MTAP* (∼60%) are present in MPNSTs ^23^. Given PRMT5’s essential roles in normal cells, targeting it with high specificity in cancer cells is particularly desirable. Thus, PRMT5 could be a potential specific target for MTAP-deficient MPNSTs. It is worth noting that a few potent PRMT5 inhibitors have been developed and tested in phase I/II clinical trials for breast cancer, lymphoma, and prostate cancer ^21,29–31^. In this study, we report that PRMT5 is highly expressed in human MPNSTs and correlates with MTAP loss. PRMT5 inhibition suppressed cancer cell growth specifically in MTAP-deficient MPNSTs, suggesting that PRMT5 represents a viable and clinically relevant therapeutic target for this type of malignancy.

## MATERIAL AND METHODS

### Immunohistochemistry (IHC) and quantification

Unstained sections of normal tissues (liver, kidney, peripheral nerve), neurofibroma, atypical neurofibroma, and MPNST were collected from the Indiana University School of Medicine. The unstained tissue on slides was dewaxed with xylene (2 × 10 min) to remove paraffin, then rehydrated with a series of diluted ethanol (100%, 95%, 80%, 50%, 5 min each) and a distilled water rinse. Then, tissues were treated with 3% H_2_O_2_ for 30min, and antigen retrieval was performed by heating slides in citrate buffer (pH=6.0) at 95°C for 30min. After blocking with 5% non-fat milk solution, anti-PRMT5 (Sigma, cat. #07-405, RRID: AB_310589, 1:500), anti-MTAP (Proteintech, cat. #11475-1-AP, RRID:AB_2147094, 1:400), and anti-H4R3me2s (Abcam, cat# ab5823, RRID:AB_10562795, 1:2000) were applied overnight. After three washes with Tris-HCl (pH 7.4) buffer solution (TBS), color staining was performed according to the instructions provided with the Vector DAB Substrate Kit and anti-rabbit secondary antibody (Vector Laboratories, cat# SK-4100, RRID:AB_2336382). DAB staining was interpreted as nuclear when the brown signal was confined within the hematoxylin-stained nuclear boundary, and as cytoplasmic when the signal was distributed outside the nucleus within the cytoplasmic region. Kidney or liver tissues were used as positive controls and normalization standards for PRMT5 and MTAP staining intensity across different batches, respectively. Immunoreactivity was graded according to the percentage of positive cells (0-100%), and staining intensity was graded as 0 (negative), 1 (weak), 2 (moderate), or 3 (strong) ^32–34^. The IHC H-score was calculated by multiplying the percentage of positive cells by the intensity (0-300)^35^. The One-way ANOVA statistical analysis was performed using GraphPad Prism 9 and RStudio.

### MPNST and neurofibroma cell lines and cell culture

MPNST cell lines (sNF96.2, RRID:CVCL_K281; sNF02.2, RRID:CVCL_K280; STS-26T, RRID:CVCL_8917; T265, RRID:CVCL_S805; ST88-14, RRID:CVCL_8916), and neurofibroma cell lines (hTERT NF1 ipNF95.6, RRID:CVCL_UI70; hTERT NF1 ipNF95.11b C, RRID:CVCL_UI67; hTERT NF1 ipnNF95.11c, RRID:CVCL_UI69; hTERT SC ipn02.3 2λ, RRID:CVCL_UI64) were acquired from American Type Culture Collection (ATCC). S462 (RRID:CVCL_1Y70), R-HT92, R-HT163, and R-HT172 MPNST cell lines were obtained from Indiana University ^36^. Dr. Christine Pratilas (Johns Hopkins University) provided JH-2-079c, JH-2-002, and JH-2-103 ^37–39^. A detailed description of the cell lines is summarized in Table S4. All cell lines were cultured in Dulbecco’s Modified Eagle’s Medium (DMEM, Corning, cat# 10-013-CV) supplemented with 10% FBS (R&D Systems, cat# S11550H) in a 5% CO_2_ environment at 37℃ ^40^. All cell cultures were confirmed to be mycoplasma-free using MycoStrip (InvivoGen, cat# rep-mysnc-50), and 1% Penicillin-streptomycin was added to some cultures as needed.

### Plasmid constructs

pLKO-TET-On shRNA plasmids for PRMT5 were efficient in human cell lines 5’-CCCATCCTCTTCCCTATTAAG-3’ (shPRMT5#1), 5’-GCCCAGTTTGAGATGC CTTAT-3’ (shPRMT5#2), and the scramble control sequence 5’-CAACAAGATGA AGAGCACCAA-3’ (shScramble) ^41^. The shRNAs were induced with doxycycline (500 ng/mL). pLix403 plasmid (Addgene, RRID:Addgene_41395) was acquired from Addgene and modified to generate pLix408 by adding EGFP via T2A self-cleavage peptide. pDONR221-hPRMT5 plasmid (DNASU:42459) was purchased from DNASU. hPRMT5 was cloned into pLix408 using Gateway LR Clonase™ II (ThermoFisher, cat# 11791020) according to the manufacturer’s instructions. The expression of hPRMT5 was induced using 500 ng/mL doxycycline. pLentiRNAGuide_001 (Addgene, RRID:Addgene_138150) and pLentiRNACRISPR_007 (Addgene, RRID:Addgene_138149) were acquired from Addgene for constructing RfxCas13d-mediated PRMT5 and MTAP knockdown ^42^. The guide RNAs for PRMT5 and MTAP knockdown are 5’-CCCCAATTTCAAGAGCCACTGCA-3’ and 5’-AGCAGTCATAATCTGTCGCCATG-3’, respectively.

### Cell lysis and Western blot

Cells were harvested in RIPA lysis buffer supplemented with 10% protease inhibitor cocktail (Millipore-Sigma, cat# 11836170001) and 10% phosphatase inhibitor cocktail (Millipore-Sigma, cat# 4906845001). Following centrifugation at 4°C, complete protein samples were prepared with 2× sample buffer and boiled at 95°C for 5min. Primary antibodies were applied accordingly: Anti-PRMT5 (Sigma, cat. #07-405, RRID: AB_310589, 1:1000), anti-β-actin (Santa Cruz Biotechnology, cat# sc-47778, RRID:AB_626632, 1:200), anti-MTAP (Proteintech, cat. #11475-1-AP, RRID:AB_2147094, 1:2000), anti-p16/Ink4a (Proteintech, cat# 10883-1-AP, RRID:AB_2078303, 1:1000), anti-SDMA (Cell Signaling Technology, cat# 13222, RRID:AB_2714013, 1:2000), anti-H4R3me2s (Abcam, cat# ab5823, RRID:AB_10562795, 1:2000), anti-phospho-Histone H2A.X(Ser139) (Millipore, cat# 05-636, RRID:AB_309864, 1:200), anti-phospho-CHK1 (Cell Signaling Technology, cat# 2348, RRID:AB_331212, 1:1000), anti-CHK1 (Cell Signaling Technology, cat# 2360, RRID:AB_2080320, 1:1000), anti-phospho-CHK2 (Cell Signaling Technology, cat# 2197, RRID:AB_2080501, 1:1000), anti-CHK2 (Cell Signaling Technology, cat# 6334, RRID:AB_11178526, 1:1000), anti-RPA32 (Santa Cruz Biotechnology, cat# sc-56770, RRID:AB_785534, 1:200), anti-Rad51 (Cell Signaling Technology, cat# 8875, RRID:AB_2721109, 1:1000), anti-Ku70 (Santa Cruz Biotechnology, cat# sc-17789, RRID:AB_628454, 1:1000) and anti-Ku80 (Cell Signaling Technology, cat# 2753, RRID:AB_2257526, 1:1000). For timelzlcourse Western blot experiments, protein levels at day 0 were used as the normalization reference for all subsequent time points after doxycyclinelzlinduced PRMT5 knockdown. For CHK1 and CHK2 phosphorylation, the ratios plzlCHK1/CHK1 and plzlCHK2/CHK2 at day 0 were used to normalize phosphorylation levels across time points.

### Stable cell line generation

TurboFect transfection reagent (Thermo Fisher, cat# R0533,) was utilized for transfection purposes. Briefly, the cocktail of 7.5μg packing plasmid, pdR8.2 (Addgene, RRID:Addgene_8455), 2.5μg envelop plasmid pVSV-G (Addgene, RRID:Addgene_8454), and 10μg plasmid of interest were prepared. The plasmid cocktail was mixed with 40 μL of Turbofect reagent, then diluted to 1mL with Turbofect growth medium. The final solution was equilibrated to room temperature for approximately 30 minutes before being added to the HEK293T (RRID: CVCL_0063) cells. In the following days, the media containing lentivirus particles was collected. After filtration through a 0.45 μm filter, the lentivirus was precipitated using a Lenti-X concentrator (Takara Bio, cat# 631232) at 4°C overnight, then centrifuged at 1,500*g* for 45 minutes. The virus pellet was resuspended in a smaller volume of fresh medium to achieve a higher titer. MPNST cells were infected overnight in complete medium containing 8 μg/mL polybrene. Following infection, cells were selected with 1-2 μg/mL puromycin for approximately 4-5 days.

### MTT assay for cell viability

Approximately 1,500-3,000 cells per well were seeded into a 96-well plate and incubated overnight at 37 °C. In the experimental wells, doxycycline or chemical inhibitors were applied on the following day. 0.5mg/mL MTT was added at 37°C for 4 hours. Then, insoluble formazan was dissolved in 100 μL DMSO. The optical density at 595nm was measured using a plate reader (Molecular Devices SpectraMax i3x Multi-Mode Microplate Reader, RRID:SCR_026346). For each experimental condition, at least three replicates were performed at each time point. The cell count was measured every 2 days for a total of 10-12 days. The area under the cell growth curves (AUC) was calculated to quantify overall proliferation. The inhibitory or promotional effects of gene knockdown or overexpression were determined by comparing the AUC of the +Dox group with that of the –Dox control group. Three biological replicates were performed for each condition.

### Colony formation assay

For each experimental condition, approximately 1000-3000 cells per well were seeded into a 6-well plate and incubated overnight at 37°C with 5% CO_2_. Cell lines were then treated with doxycycline or inhibitors for another 6-14 days, until visible colonies formed in the untreated control groups. Fresh medium containing doxycycline (500 ng/mL) was replaced every 3 days. Colonies were visualized using 0.05% crystal violet in 6% glutaraldehyde. Colonies with diameters >0.2 μm were counted.

### Flow cytometry

For cell cycle analysis, cells were prepared and cultured with 500 ng/mL doxycycline for the expected time periods, then harvested and fixed in 95% Ethanol overnight. After a blocking step with 4% BSA, anti-phospho-histone 3 (Santa Cruz Biotechnology, cat# sc-8656-R, RRID:AB_653256, 1:200) was applied overnight. The secondary antibody DyLight 488 (Thermo Fisher Scientific, cat# 35552, RRID:AB_844398, 1:250) was used, followed by propidium iodide DNA content staining (50μg/mL) at 4 °C overnight. Samples were then ready for flow cytometry. Flow cytometry was performed on Attune NxT (Thermo Fisher Attune Nxt Flow Cytometer, RRID:SCR_019590), and results were analyzed with FlowJo (Version 10.8.1, RRID:SCR_008520). For doxycyclinelzlinduced PRMT5 knockdown, the celllzlcycle distribution at day 0 served as the control. Subpopulation percentages at the subsequent time points were compared to the daylzl0 distribution. Three biological replicates were analyzed for each time point.

For cell apoptosis analysis, after 12 days of treatment with 500 ng/mL doxycycline, cells were collected and washed in cold phosphate-buffered saline (PBS) on ice. Samples were diluted to 1 x 10^6^ cells/mL with 1 x Annexin-binding solution (10mM HEPES, 140mM NaCl, 2.5mM CaCl_2_, pH=7.4). 5 μL Alexa Fluor 488 Annexin V (Invitrogen, cat# V13241) and 1 μg/mL propidium iodide were applied to 100 μL of cell suspension. Samples were incubated at room temperature for 15 minutes, then gently mixed with 400 μL 1x Annexin-binding solution.

### Cell cycle analysis using Fucci constructs

The plasmid pBOB-EF1-FastFUCCI (Addgene, RRID:Addgene_86849) was utilized to generate a stable cell line by lentivirus transduction as described above and selected by 2 μg/mL puromycin for 5 days. The fluorescence signals (RFP and GFP) were monitored under the inverted fluorescent microscope (Zeiss Axio Vert series Axiovert 200 inverted microscope, RRID:SCR_020915). For cell cycle stage evaluation, cells were first synchronized by overnight serum starvation, followed by 3 days of treatment with 0.01 μM JNJ-64619178. A minimum of five randomly selected fields per sample were imaged under the confocal microscope (Nikon ECLIPSE Ti2 inverted microscope, RRID:SCR_021068) equipped with a spinning-disk unit (CrestOptics CICERO). The images were captured in NIS-Elements Advanced Research (RRID:SCR_027181). RFP- and GFP-positive cell percentages were counted to assess cell cycle progression. The average percentages of each cell cycle stage were calculated for statistical analysis. A multiple t-test was performed in GraphPad Prism.

### Comet assay

The alkaline comet assay was performed to evaluate both DNA single-strand and double-strand breaks ^43^. After doxycycline treatment for the indicated period, cells were trypsinized, washed twice with PBS, and resuspended in PBS. For each experimental condition, 8,000-10,000 cells in 400 μL PBS were gently mixed with 1.2 mL 1% low-temperature agarose solution at 40°C. Then, the mixed solution was placed on a 75 x 25 mm microscope slide (Corning, cat# CLS294875X25), and the agarose was allowed to solidify at room temperature. The slides containing the gels were submerged in alkaline lysis buffer (1% SDS, 100 mM Na2EDTA, 1.2 M NaCl, 0.26 M NaOH, pH > 13) at 4°C overnight. After lysis, slides were rinsed twice for 20min and equilibrated in running buffer (0.03M NaOH, 2mM Na_2_EDTA, pH=12.3) for 30min at room temperature, followed by electrophoresis at 20V for 30min. DNA was visualized by incubating in a 2.5 μg/mL propidium iodide solution for 20 minutes, followed by three rinses with distilled water. Images were captured using a fluorescence microscope (Zeiss Axio Imager 2, RRID:SCR_018876) equipped with a color camera (Zeiss AxioCam MRm). Tail moment indicating DNA breaks was automatically calculated by OpenComet (RRID:SCR_021826) based on tail length and staining intensity ^43^. Tail moments measured at the various time points after doxycyclinelzlinduced PRMT5 knockdown were compared to the tail moments at day 0. At least 50 cells were analyzed per time point. At least 50 cells were counted for each sample.

### Immunofluorescence staining of cultured cells

Cells were seeded on the sterile coverslip in the culture disk overnight. 500 ng/mL doxycycline was administered. On the day indicated, cells were washed 3 times with PBS for 5 minutes each, then fixed with 4% PFA overnight at 4°C, followed by 15 minutes of PBS washing at room temperature. Then, cells were permeabilized by incubation in PBST (0.3% Triton X-100) at room temperature for 30 minutes. After blocking with 4% BSA for one hour at room temperature, anti-PRMT5 (Sigma, cat. #07-405, RRID: AB_310589, 1:500), anti-phospho-Histone H2A.X(Ser139) (Millipore, cat# 05-636, RRID:AB_309864, 1:200), and anti-RPA32 (Santa Cruz Biotechnology, cat# sc-56770, RRID:AB_785534, 1:200) were applied and incubated at 4℃ overnight. Slides were washed twice with PBST (0.3% Triton-X 100) followed by incubation of the secondary antibody DyLight 488 (Thermo Fisher Scientific, cat# 35552, RRID:AB_844398, 1:250) or Alexa Fluor 647 (Jackson ImmunoResearch Labs, cat# 111-605-003, RRID:AB_2338072, 1:250) at room temperature for 2h. Coverslips were mounted on the slides after washing, and the signals were detected under the confocal microscope (Nikon ECLIPSE Ti2 inverted microscope, RRID:SCR_021068). For each sample, images were acquired under the identical microscope settings in NIS-Elements Advanced Research (RRID:SCR_027181). All non-overlapping fields were imaged at low magnification (40x) sequentially to capture the entire cell area. Then, high-magnification (200x) images were acquired in the low-magnification (40x) fields that included visible cells. At least five high-magnification (200x) images per sample were analyzed. An unpaired t-test was used for statistical analysis.

### IC_50_ assay and cell treatment of chemical inhibitors

The PRMT5-specific chemical inhibitors JNJ-64619178 (CAS No. 2086772-26-9), MRTX1719 (CAS No. 2630904-45-7), GSK3326595 (CAS No. 1616392-22-3), and PF-03639999 (CAS No. 2159123-14-3) were applied to the selected MPNST cancer cells. All four PRMT5 inhibitors were purchased from Chempedak. A 10 mM stock was prepared and applied to the cell lines at the indicated concentrations. JNJ-64619178, MRTX1719, and PF-03639999 were dissolved in sterile water, and GSK3326595 was dissolved in DMSO. To evaluate the sensitivity of the cell lines to the inhibitors and to select appropriate treatment concentrations for MTAP-deficient MPNST cell lines, dose-response curves with half-maximal inhibitory concentration (IC_50_) estimates were generated. Each inhibitor was prepared in a series of dilutions (10-point) spanning a broad concentration range with the expected activity. Cells were treated for 5 days, then an MTT assay was performed to measure cell numbers. Three replicates were performed for each group. Drug response curves were generated in GraphPad Prism 9 (RRID:SCR_002798) using the log[inhibitor] vs normalized response (variable slope) equation model based on the 10 observed points, and IC_50_ values were estimated ^44^. Following the IC_50_ assay, appropriate concentrations of the PRMT5 inhibitors were selected to validate specific inhibitory effects in comparison between the MTAP-deficient and MTAP-proficient MPNST cell lines. Inhibitory effects were quantified by calculating and comparing the area under the curve (AUC) for the treatment conditions with that of the untreated groups ^45^.

### Combination treatment

Doxorubicin (CAS No.25316-40-9), gemcitabine (CAS No. 95058-81-4), and 5-FU (CAS No. 51-21-8) were purchased from Sigma-Aldrich. Cell lines were seeded at 500-1000 cells per well in 96- and 6-well plates for overnight incubation and pretreated for 5 days with 500 ng/mL doxycycline. Similarly, 10 gradient concentrations of doxorubicin/gentamicin/5-FU were applied for 5 days, followed by MTT assay for cell growth or crystal violet staining for colony formation. IC_50_ analysis was performed using MTT assay results, as described previously, to evaluate the effects of chemotherapy on cell lines with and without PRMT5 knockdown. For combination drug treatment, JNJ-64619178 and MRTX1719 were tested at five concentrations, along with one untreated control, in combination with four doxorubicin concentrations and one untreated control, and applied to the cell lines for 5 days. Cell number was evaluated using the MTT assay. The synergistic effect parameters were analyzed by Combenefit software (RRID:SCR_027410) according to the Loewe model ^46–48^. Three replicates of each combination treatment group were performed.

### β-galactosidase staining for cellular senescence

Cells were seeded in 6-well plates and allowed to reach 70–80% confluency. On the 12^th^ day, cells were washed twice with PBS and fixed with 4% PFA for 15 minutes at room temperature. After fixation, cells were washed three times with PBS and incubated overnight at 37°C (in a dry incubator without CO₂) with freshly prepared staining solution containing 1 mg/mL X-gal (Thermo Fisher, cat# B1690), 40mM citric acid/sodium phosphate buffer (pH 6.0), 5mM potassium ferrocyanide, 5mM potassium ferricyanide, and 150 mM NaCl. Following incubation, stained cells were washed with PBS, and senescent cells were identified by their blue staining under bright-field microscopy (Zeiss Axio Vert series Axiovert 200 inverted microscope, RRID:SCR_020915). The non-overlapping fields were imaged at low magnification (60x) sequentially to capture the entire cell area. Then, high-magnification (100x) images were acquired in the low-magnification (60x) fields that included visible cells. At least five high-magnification (100x) images per sample were analyzed. Cells showing distinct diffuse blue cytoplasmic staining covering >50% of the cell area were counted as positive, whereas unstained or faintly stained cells were considered negative. Results are expressed as the percentage of SA-β-gal-positive cells relative to the total number of cells per field.

### Analysis of public datasets

Transcriptomic data from whole-exome sequencing of nerve sheath tumor samples with their matched normals, including PRMT5 and MTAP mRNA expression, were downloaded from cBioPortal (https://www.cbioportal.org/, RRID:SCR_014555) ^49,50^. Data was analyzed and plotted by using GraphPad Prism 9 (RRID:SCR_002798) and RStudio (RRID:SCR_000432). One-way ANOVA and simple linear regression statistical analysis were performed. Kaplan-Meier survival analysis of sarcoma patients stratified by PRMT5 mRNA levels (TPM) was generated on GEPIA (http://gepia.cancer-pku.cn/, RRID:SCR_018294) ^51^. The highest and lowest quartiles (cutoff 75% and 25%) of PRMT5 mRNA levels (TPM) among all sarcoma patients defined high and low PRMT5 expression groups, respectively. Log-rank statistical analysis was used. Human RPA trimeric core structure (RCSB PDB ID: 8RK2), including all three subunits RPA70, RPA32, and RPA14, was obtained from the Protein Data Bank (https://www.rcsb.org/, RRID:SCR_012820) ^52,53^. The predicted structure for human RPA32 (UniProt ID: P15927) was downloaded from the AlphaFold Protein Structure Database (https://www.alphafold.ebi.ac.uk, RRID:SCR_023662) ^54^. Three top-ranking potential PRMT5 di-methylation sites on RPA32 were predicted by AlphaFold2 (RRID:SCR_025454) based on a consensus amino acid motif (RG-rich domain) containing PRMT5-mediated di-methylation of arginine and on accessibility (highly disordered domain) ^55^.

## RESULTS

### Increased PRMT5 expression in MTAP-deficient MPNSTs

Although MTAP loss and PRMT5 upregulation have been reported in some genomic studies, confirmation of these findings by protein expression in human MPNSTs has been lacking. To address this gap, immunohistochemistry was used. Approximately half of MPNST cases are sporadic, and the other half are associated with NF1. The NF1-related MPNSTs typically exhibit a stereotyped progression, characterized by benign neurofibroma, premalignant atypical neurofibroma, and low- and high-grade MPNSTs. To gain a comprehensive understanding of PRMT5 and MTAP expression patterns, we collected samples from 42 neurofibromas (NF), 5 atypical neurofibromatous neoplasm with uncertain biologic potential (ANNUBP), 33 MPNSTs (both NF1 and non-NF1), and 31 peripheral nerve tissues as the normal control.

MTAP and PRMT5 expression are highly variable across our samples, not only in levels but also in cellular localization (cytoplasmic and nuclear). MTAP was predominantly found in the cytoplasm of the normal peripheral nerve, while nuclear MTAP was more prevalent in the neurofibroma and MPNSTs in our study (Fig. 1A-E). To avoid confusion, we focused only on cytoplasmic MTAP in this study, as it is physiologically involved in the methionine salvage pathway. In contrast, the functions of nuclear MTAP are less well characterized. Overall, 85% (28/33) MPNSTs, all 5 ANNUBPs, and 80% neurofibromas (34/42) decreased or completely lost MTAP expression compared to the normal peripheral nerves (Fig. 1A-E). Among MPNST cases, 50% (3/6) low-grade MPNSTs presented as abundant MTAP as the normal peripheral nerve, but 93% (25/27) high-grade MPNSTs had no or less MTAP (Fig. 1K). These results demonstrate that MTAP can be lost in MPNSTs, consistent with the reported frequent MTAP deletions in MPNSTs associated with *CDKN2A* loss^1,2^.

**Figure 1.**
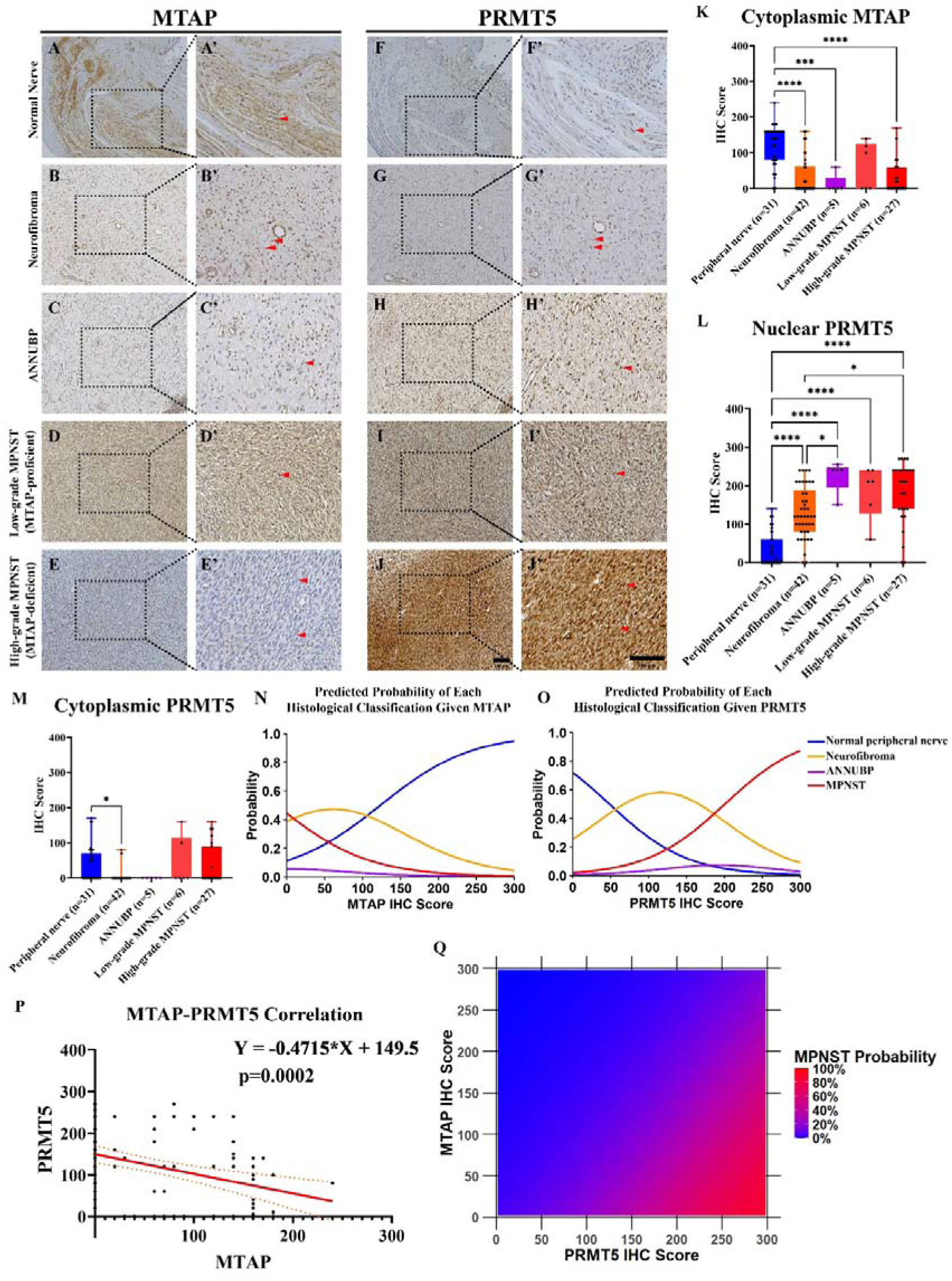
MTAP loss and increased PRMT5 expression during MPNST tumorigenesis. **A-J.** Immunostaining analysis of MTAP (left) and PRMT5 (right) in the normal peripheral nerve (**A**, **F**), neurofibroma (**B**, **G**), ANNUBP(**C**, **H**), and MPNST (**D**, **E**, **I**, **J**). The representative images were taken at the 100x (**A**-**J**) and 200x (**A’**-**J’**) magnifications. Red arrows indicate the nuclei of peripheral nerve or tumor cells. **K**. Quantification of MTAP expression. Only cytoplasmic MTAP staining was counted. The area of immunoreactivity was graded from 0% to100%, and the intensity of staining was categorized as 0, negative; 1, weak; 2, moderate; and 3, strong. IHC H-score was calculated by multiplying the extent and intensity (0-300). ***p<0.001, ****p<0.0001, ns. Nonsignificant. **L**. Quantification of nuclear PRMT5 staining. *p<0.05, ****p<0.0001. ns. Nonsignificant. **M**. Quantification of cytoplasmic PRMT5 staining. *p<0.05. ns. Nonsignificant. **N**. Predicted probability of the physiological and pathological states (normal peripheral nerve, neurofibroma, ANNUBP, and MPNST) with cytoplasmic MTAP expression using the ordinal logistic regression model. **O**. The predicted probability of the physiological and pathological states (normal peripheral nerve, neurofibroma, atypical neurofibroma, and MPNST) with nuclear PRMT5 expression using the ordinal logistic regression model. **P**. General linear correlation between cytoplastic MTAP and nuclear PRMT5 expression in the normal peripheral nerve, neurofibroma, atypical neurofibroma, and MPNST. **Q**. The estimated probability of MPNST with cytoplasmic MTAP and nuclear PRMT5 expression by using the ordinal logistic regression model.

Next, we examined PRMT5 protein levels and found increased PRMT5 levels in neurofibroma (NF), ANNUBP, and MPNST compared with normal peripheral nerve (Fig. 1F-J). Similar to MTAP, PRMT5 was also found in both the cytoplasm and the nucleus in our study, consistent with its diverse functions. Therefore, we analyzed PRMT5 in both cellular locations in these samples. A low level of nuclear PRMT5 was detected in the normal peripheral nerves. In contrast, strong nuclear PRMT5 expressions were observed in neurofibroma, ANNUBP, and MPNSTs (Fig. 1L). Interestingly, there was no significant difference in cytoplasmic PRMT5 between normal peripheral nerve and tumors (Fig. 1M), suggesting that PRMT5 dependency in MPNST was associated with nuclear rather than cytoplasmic activities. We also explored whole-exome sequencing data of neurofibromas and MPNSTs on cBioPortal and found consistent increases in PRMT5 mRNA levels in MPNSTs compared to the benign tumors (Fig. S1A). For MTAP mRNA, there was no significant difference between benign tumors and MPNST, consistent with our MTAP IHC results (Fig. S1B). The MTAP data implied that MTAP loss could occur in benign neurofibroma at a relatively early stage of tumorigenesis. Moreover, we calculated and found a decreased MTAP/PRMT5 mRNA ratio in MPNSTs, suggesting consistent changes of MTAP and PRMT5 patterns with those in IHC results (Fig. S1C).

To understand the correlational changes between MTAP and PRMT5 from normal nerves to MPNSTs, we performed ordinal logistic regression analyses of both proteins across the four categories. We found a significant predicted trend indicating that a normal peripheral nerve was likely to present high MTAP expression. In contrast, the probability of pathological NF, ANNUBP, and MPNST increased with decreasing or absent MTAP (Figure 1N). Moreover, high-grade MPNSTs exhibited more prevalent MTAP loss than low-grade MPNSTs (Table S1). Correspondingly, low nuclear PRMT5 expression was the most prevalent in the normal peripheral nerve. However, higher PRMT5 expression was associated with a higher probability of MPNST and atypical neurofibroma; neurofibroma was most likely to have moderately increased PRMT5 (Fig. 1O). Based on our hypothesis, the increase in PRMT5 may reflect physiological compensation for MTAP loss. Thus, we explored the association between PRMT5 and MTAP expression and found a highly significant, negative correlation in NFs, ANNUBPs, and MPNSTs (Fig. 1P). The ordinal logistic regression model also suggested that low MTAP and high PRMT5 were strongly associated with the high probability of MPNST (Fig. 1Q). Our public dataset analysis of MTAP and PRMT5 whole-exome sequencing data, although not significant, also implied a correlation of decreased MTAP and increased PRMT5 mRNA levels across benign nerve sheath tumors and MPNSTs (Fig. S1D). Overall, our results indicate that cytoplasmic MTAP loss and nuclear PRMT5 increase are hallmarks of MPNST tumorigenesis.

### Increased PRMT5 function in MTAP-deficient MPNSTs

Increased protein expression does not always reflect increased activity, as additional mutations may occur. To further validate the increased dependence on PRMT5 activity in MTAP-lost tumors, the PRMT5 substrate and activity marker, H4R3me2s (histone 4 arginine 3 symmetrical dimethylation), was evaluated in ANNUBPs and MPNSTs. MTAP-proficient MPNST cases showed relatively lower H4R3me2s staining than MTAP-deficient tumors. In contrast, MTAP-deficient ANNUBP and MPNST had stronger PRMT5 and H4R3me2s (Fig. 2A-I). Thus, increased PRMT5 expression matched higher PRMT5 activity in the majority of samples. We aligned all ANNUBP and MPNST cases with MTAP, PRMT5, and H4R3me2s levels based on MTAP levels. We found that strong PRMT5 expression and activity were enriched in the high-grade cases (Fig. 2J). Consistently, our statistical analysis also revealed that increased PRMT5 activity was enriched in the MTAP-deficient and high-grade MPNSTs (Fig. 2K-L). However, it is worth noting that among the MTAP-deficient cases (3 out of 34), low PRMT5 expression was also occasionally observed (Fig. 2L). This phenomenon may be caused by PRMT1 or PRMT9 compensating for PRMT5, or simply by a hypermorphic PRMT5 mutation in these samples.

**Figure 2.**
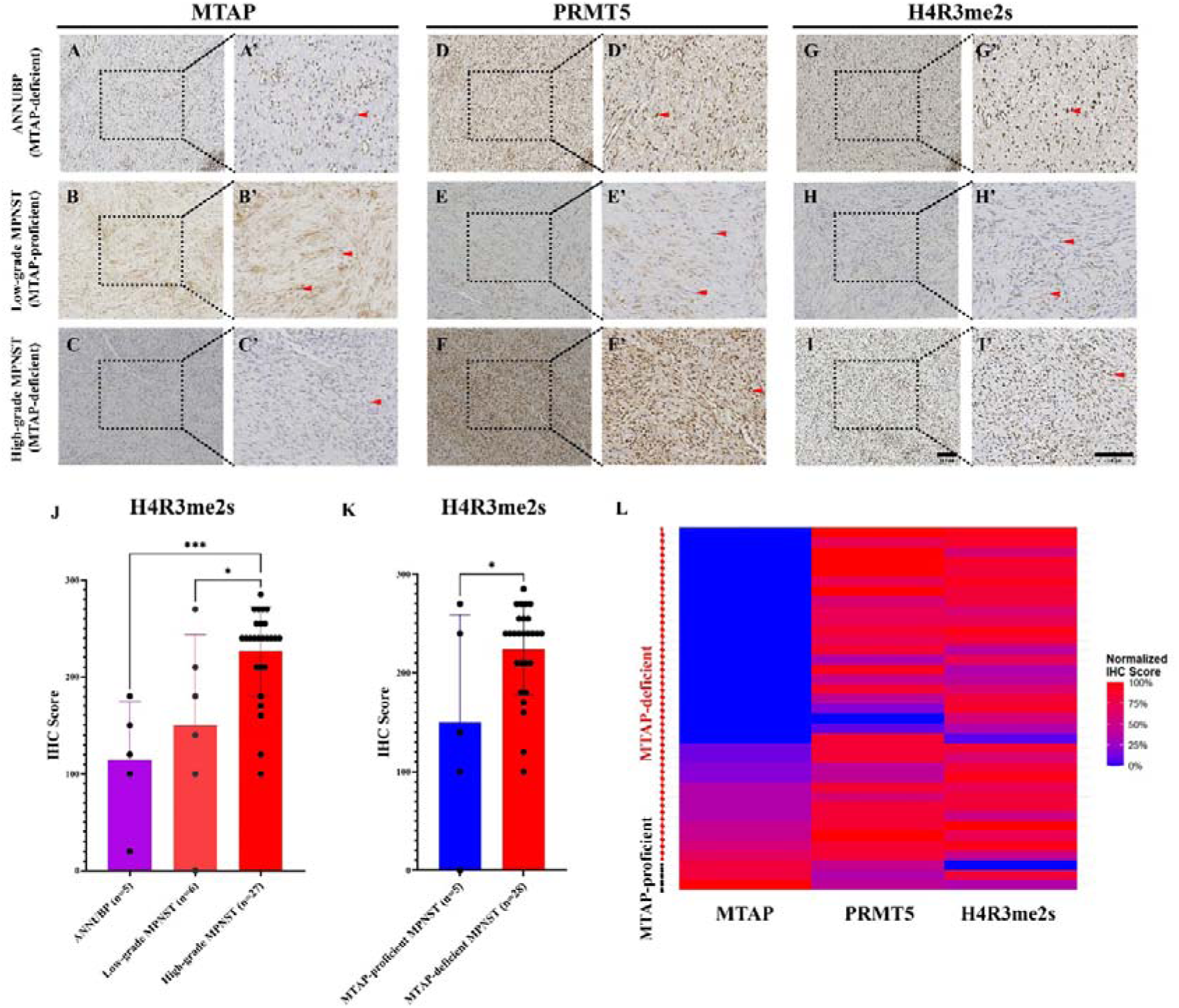
MTAP loss was correlated with increased PRMT5 activity. **A**-**I**. Immunostaining analysis of MTAP, PRMT5, and H4R3me2s on adjacent tissue sections of the ANNUBP, low-grade, and high-grade MPNST. The representative images were taken at 100x (**A**-**I**) and 200x (**A’**-**I’**) magnifications. Red arrows labeled the nuclei of tumor cells. **J**. Quantification of the H4R3me2s expression in the ANNUBP, low-grade, and high-grade MPNST. *p<0.05, ***p<0.001. **K**. The quantification of the H4R3me2s expression in the MTAP-proficient and MTAP-deficient ANNUBP and MPNST. *p<0.05. **L**. Heatmap showing the paired PRMT5 and H4R3me2s with the MTAP levels in the same atypical neurofibroma and MPNST.

Next, we examined the relationship between patient survival and PRMT5 expression levels. Among all parameters, patients with metastatic or recurrent MPNSTs had significantly shorter survival times than those without metastasis or relapse, as expected (Tables S2-3). There was a trend toward increased PRMT5 expression in metastatic and recurrent cases, although the differences were not statistically significant, possibly due to the limited number of cases and tissue samples. This suggests a crucial requirement for high PRMT5 activity in these advanced MPNSTs. Furthermore, survival analysis of the sarcoma patients in the TCGA dataset showed poorer survival in patients with high PRMT5 expression (Fig. S1E), consistent with our MPNST clinical results.

### PRMT5 knockdown inhibited the cell growth of MTAP-deficient MPNST cells

Following confirmation that MTAP is frequently lost and PRMT5 expression is upregulated in human NF and MPNST samples, we evaluated MTAP and PRMT5 expression in 12 MPNST and 6 transformed benign neurofibroma/schwannoma cell lines (Table S4). Using the benign NF cell line as a reference, ipNF95.11b C, which had the highest MTAP expression among the benign transformed cell lines, we identified seven MPNST and three benign transformed cell lines with significantly low to undetectable MTAP expression and five MPNST cell lines with proficient MTAP expression (Fig. 3A-B, Fig. S2A). Low MTAP protein expression is not always associated with *CDKN2A* loss (Fig. 3A), as shown in a recent study of peritoneal mesothelioma ^56^. PRMT5 levels are readily detectable in all MPNST cell lines and are similar to those in the neurofibroma line; however, the PRMT5/MTAP ratio is evident in six MTAP-deficient cell lines (Fig. 3B).

**Figure 3.**
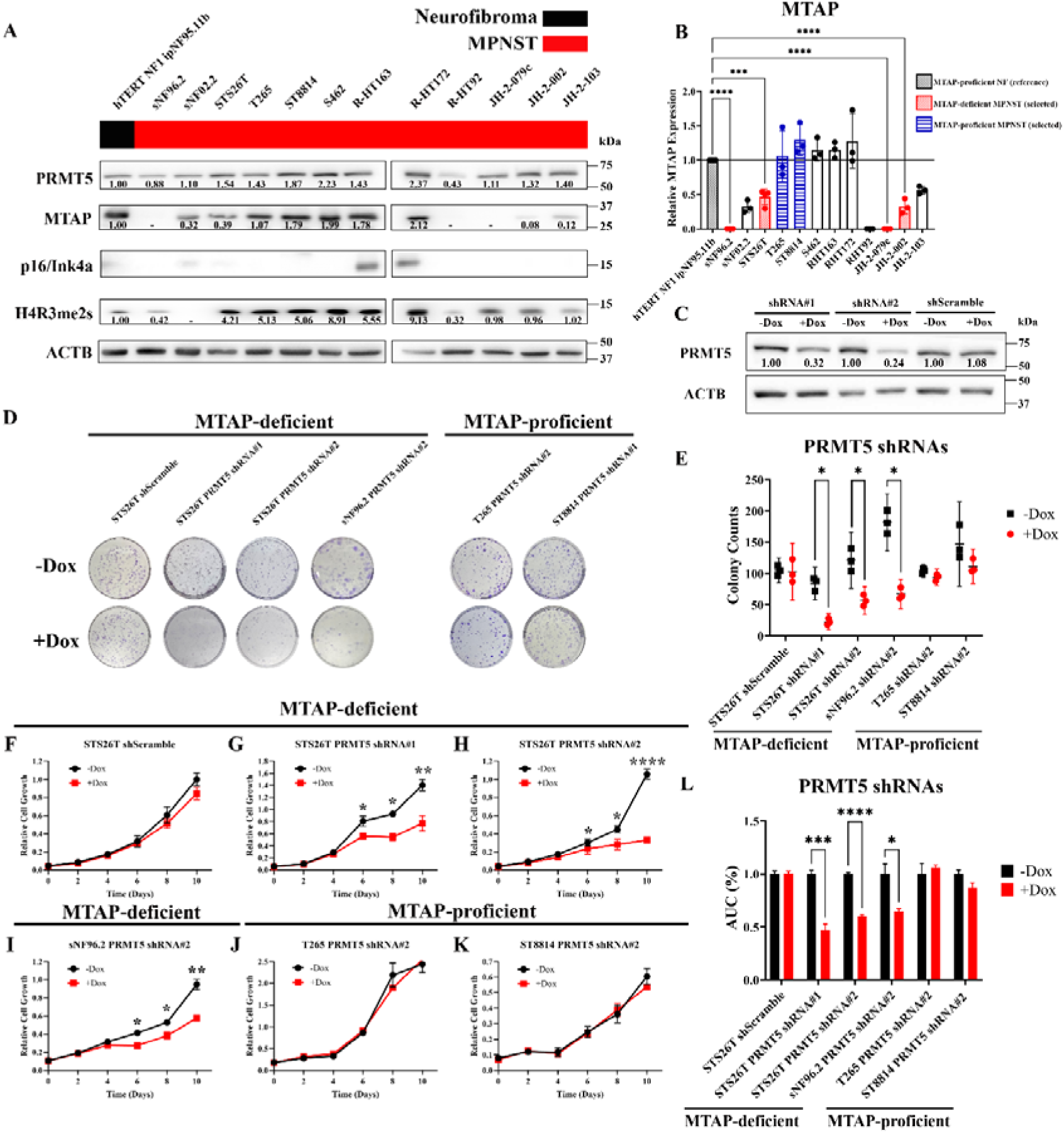
PRMT5 Knockdown Inhibited growth of MTAP-deficient MPNST Cancer Cells. **A**. Western blot analysis of the endogenous PRMT5, MTAP, H4R3me2s (PRMT5 activity marker), and p16/Ink4a (CDKN2A) in the MPNST cell line collection with the control transformed benign neurofibroma cell line hTERT NF1 ipNF95.11b C. ACTB was used as the loading control. **B**. Relative MTAP levels were quantified compared to the control cell line hTERT NF1 ipNF95.11b C. The quantification of the MTAP endogenous levels of the MPNST cell lines compared to the control cell line ipNF95.11b C. Three independent replicates for each cell line. *p<0.05, ****p<0.0001. Red: deficient MTAP; Blue: proficient MTAP. **C**. Efficiencies of the 2 PRMT5 shRNAs (#1 and #2) and 1 scramble shRNA as the control by Western blot analysis. The representative gels showed the PRMT5 expression levels in PRMT5 shRNA or shScramble STS-26T cell lines with or without 5-day Dox+ (500 ng/mL) treatment. The efficiencies (numbers at the bottom of the upper gel) of the shRNAs were quantified by comparing PRMT5 expression levels of the Dox+ to the Dox-group in the cell lines. Dox, doxycycline. **D**-**E**. Effects of the PRMT5 shRNAs (#1 and #2) or the control scramble RNA on the colony formation of the MPNST cell lines. Dox treatment was administered for 8-12 days. The colony was visualized via crystal violet staining. The colonies with diameters> 0.2 μm were counted. Three replicates were performed for each group. *p<0.05, **p<0.01, ***p<0.001. ns. Nonsignificant. **F**-**K**. Effects of the PRMT5 knockdown by shRNAs (#1 and #2) or the control scramble RNA on the overall cell growth of the MPNST cell lines by MTT assay. Cell counts of the control group (black: Dox-) and the PRMT5 knockdown group (red: Dox+) were evaluated every 2 days for a total of 10 days using the MTT assay. OD595 values were measured for 3 replicates in each condition. *p<0.05, **p<0.01, ****p<0.0001. **L**. Quantification of PRMT5 knockdown effects by comparing the area under the curve (AUC) of the Dox-control group with that of the Dox+ treatment groups.

To examine whether PRMT5 is essential for the low MTAP cells, we selected MTAP-deficient (sNF96.2 and STS-26T) and MTAP-proficient MPNST (T265 and ST-8814) cell lines. First, PRMT5 was knocked down using previously validated shRNAs ^41^ (Fig. 3C), and cell growth was examined using colony formation and MTT assays. Growth of MTAP-deficient MPNST cell lines was significantly reduced compared to that of MTAP-proficient cell lines by both assays (Fig. 3D-L). This growth-inhibitory effect of PRMT5 genetic inhibition was replicated using the doxycycline-inducible RfxCas13-CRISPR system (Fig. S2). Thus, our results support targeting PRMT5 for MTAP-deficient MPNSTs. Notably, we found a time lag between the cell-growth-inhibitory effects of PRMT5 genetic knockdown and the decrease in PRMT5 expression. PRMT5 expression was significantly reduced by shRNA and RfxCas13d on the 3^rd^ and 2^nd^ days, respectively, and reached the lowest levels on the 5^th^ and 3^rd^ days, respectively (Fig. S3). In contrast, the decrease in cell growth was noticeable on the 6^th^ day (Fig. 3G-I; Fig. S2B). Interestingly, PRMT5 knockdown resulted in significantly reduced cell growth in MTAP-deficient MPNST cell lines; however, complete abrogation of cell growth was not observed (Fig. 3F-L; Fig. S2B).

### MTAP-deficient MPNST cells are sensitive to PRMT5 inhibitors

To further evaluate the therapeutic potential of PRMT5 inhibition in MTAP-deficient MPNSTs, we selected four PRMT5 chemical inhibitors: JNJ-64619178, MRTX1719, PF-0693999, GSK3326595. Generally, PRMT5 inhibitors are categorized into two categories based on their mechanisms of action. JNJ-64619178, PF-0693999, and GSK3326595 competitively bind to the SAM-binding or substrate pockets of PRMT5 ^31,57,58^. The second type of inhibitors, such as MRTX1719, was designed to stabilize the PRMT5-MTA complex ^59^. Thus, this type of inhibitor is better suited to the MTAP-deficient condition. First, we confirmed the efficiency of these four inhibitors by examining H4R3me2s, as a readout of PRMT5 methylation function. As expected, all four inhibitors are effectively blocking PRMT5 activity in the MTAP-deficient MPNST cell line STS-26T in a dose-dependent manner (Fig. 4A).

**Figure 4.**
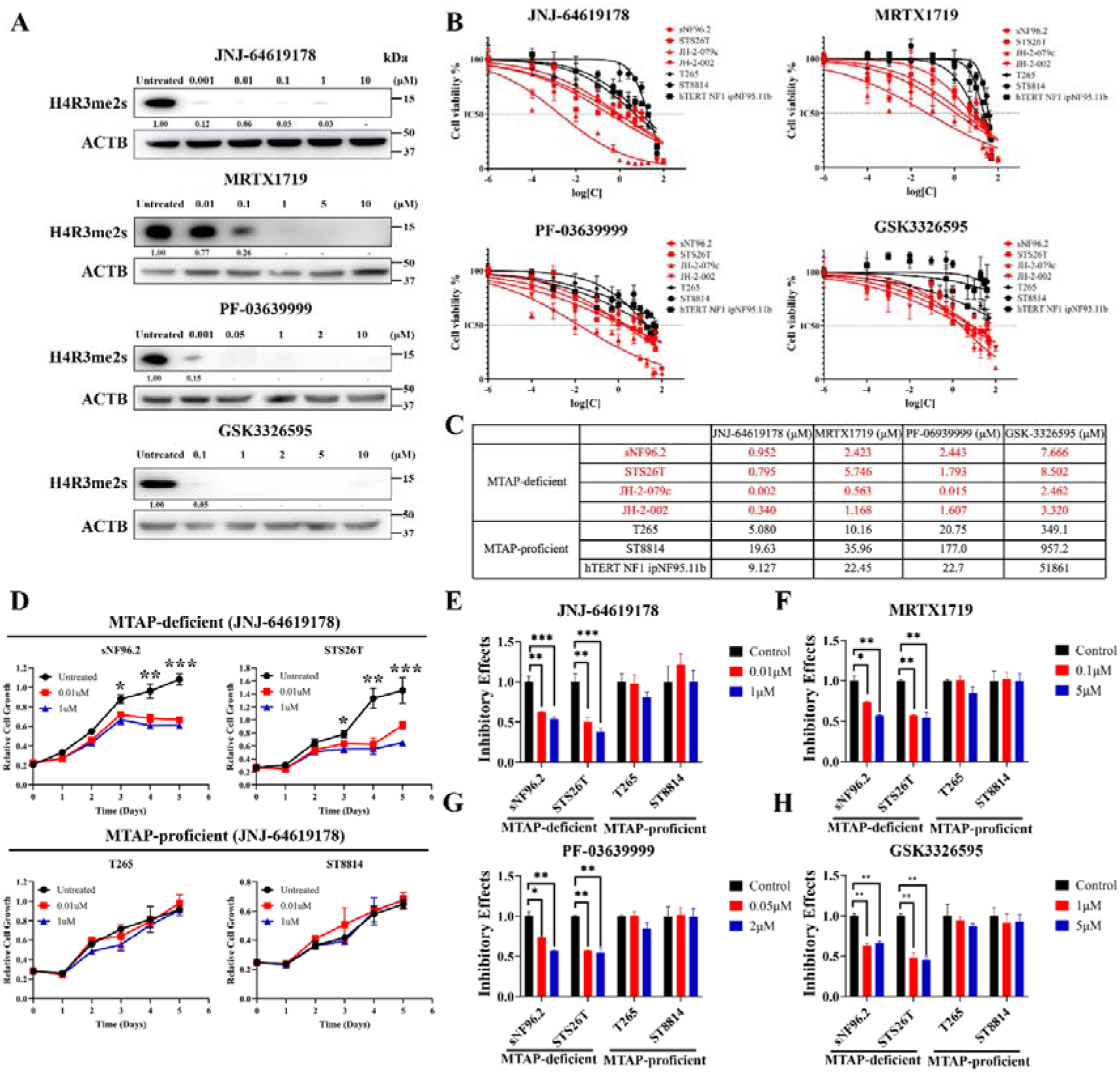
MTAP-deficient MPNST cell lines were sensitive to the potent PRMT5 inhibitors. **A**. Efficiency of the four PRMT5 inhibitors in the MTAP-deficient MPNST cell line STS-26T by Western blot analysis. The intensity of H4R3me2s was quantified relative to the untreated control. **B**. IC_50_ analysis of 4 inhibitors. Drug concentration response curves were generated based on the 10 observed points. Each observation, consisting of three replicates, represented the survival percentage at a single concentration of each inhibitor. The model of log(concentration)-response with four parameters was used. **C**. Estimated IC_50_ values based on the drug concentration response curves for each MPNST cell line. **D**. Effects of the PRMT5 inhibitors JNJ-64619178 on the overall cell growth of the MPNST cell lines by MTT assay. Cell counts for the control (black), untreated group, and PRMT5 inhibitor treatment groups at two relatively low concentrations (red and blue) were evaluated daily for a total of 5 days. OD595 values were measured for 3 replicates in each condition. *p<0.05, **p<0.01, ***p<0.001. **E**-**H**. Overall inhibitory effects of four PRMT5 inhibitors across cell lines were calculated by comparing the area under the curve (AUC) of the untreated or DMSO control group with that of the treatment groups. The initial cell counts measured on day 0 served as the baselines for the calculation. *p<0.05, **p<0.01, ***p<0.001. ns. Nonsignificant.

Next, we determined the IC_50_ for each inhibitor in each cell line, given their diverse genetic backgrounds. Overall, we found that MTAP-deficient MPNST cells (JH-2-079c, JH-2-002, sNF96.2, and STS-26T, red curves) were more sensitive to PRMT5 chemical inhibitors compared to the MTAP-proficient MPNST or neurofibroma cell lines (T265, ST-8814, and hTERT NF1 ipNF95.11b C, black curves) (Fig. 4B). IC_50_ values of the four MTAP-deficient MPNST cell lines were consistently lower than 10μM and 10-fold lower, at least, than those of the MTAP-proficient MPNST or neurofibroma cells (Fig. 4C). Then, we selected two relatively low concentrations for all four inhibitors based on IC_50_ assay and previous reports, and evaluated their inhibitory effects on overall cell growth in the two pairs of MTAP-deficient (sNF96.2, STS-26T) and MTAP-proficient (T265, ST-8814) cell lines. Consistently, the PRMT5 inhibitors at the selected concentrations effectively inhibited the growth of MTAP-deficient MPNST cells but did not affect MTAP-proficient MPNST cells (Fig. 4D-H). Our results suggest that patients with low MTAP and high PRMT5 might have a better response to PRMT5 inhibition.

### The sensitivity to PRMT5 inhibitors was dependent on MTAP protein expression levels

To further confirm that sensitivity to PRMT5 inhibition depends on MTAP expression levels, we knocked down MTAP in MTAP-proficient tumor cell lines, neurofibroma ipNF95.11b C and MPNST ST-8814, using the RfxCas13d system. ST-8814 was chosen based on its lack of response to PRMT5 inhibitors, as shown in Figure 4. With the tetracycline-inducible system, PRMT5 expression gradually and significantly increased following MTAP knockdown in the MTAP-proficient ipNF95.11b C and ST-8814 cell lines (Fig. 5A-B). Underscoring the cell’s dependence on expression of PRMT5 in the absence of MTAP, doxycycline withdrawal resulted in restoration of MTAP expression and return of PRMT5 to basal levels (Fig. 5A-B). This reverse correlation between PRMT5 and MTAP is consistent with our IHC results of human MPNST samples. Upon depletion of MTAP in an MTAP-proficient cell, the sensitivity of ipNF95.11b C and ST-8814 to the PRMT5 inhibitors, JNJ-64619178 and MRTX1719, was examined. Indeed, the two cell lines became sensitive to the inhibitors, like the MTAP-deficient sNF96.2 and STS-26T cells (Fig. 5C-H). Thus, reduced or lost MTAP contributes to the sensitivity of PRMT5 inhibition in MPNST cells. It is worth noting that MTAP knockdown has a greater effect on the proliferative capacity of the neurofibroma than on the tumor line, suggesting that tumor cells have evolved mechanisms to circumvent MTAP deletion, at least in part through PRMT5.

**Figure 5.**
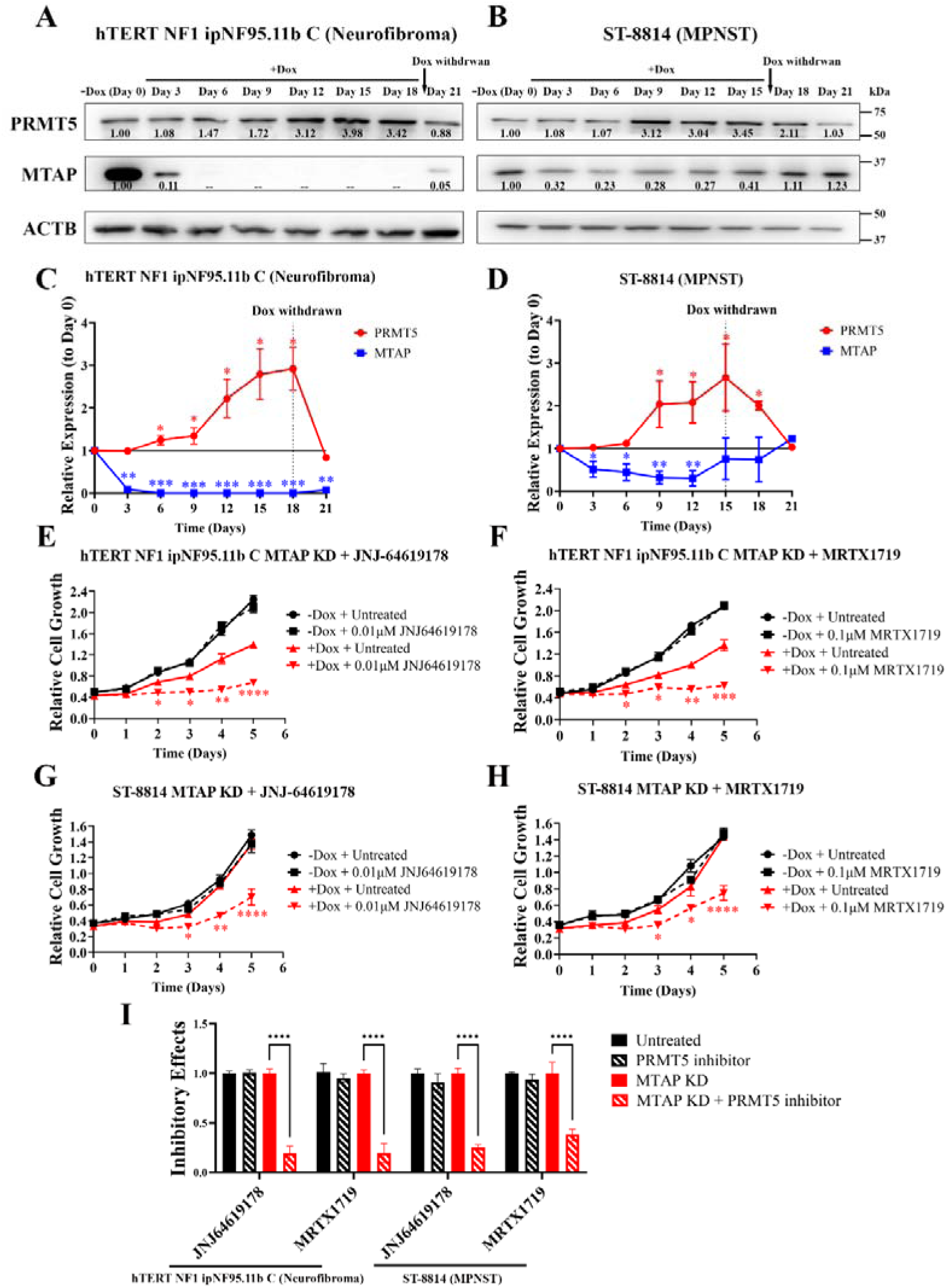
MTAP knockdown led to increased PRMT5 expression and increased vulnerability to PRMT5 inhibition in MTAP-proficient MPNST cells. **A-B**. Time course western blot analysis of PRMT5 expression changes upon MTAP knockdown mediated by Cas13d gRNAs in the MTAP-proficient neurofibroma cell line hTERT NF1 ipNF95.11b C and the MPNST cell line, ST-8814. Dox, doxycycline. **C-D**. Quantification of MTAP and PRMT5. Expression levels on day 0 (-Dox) were used as the normalization control. **E-H**. Inhibitory effects of the PRMT5 inhibitors JNJ-64619178 and MRTX1719 in the MTAP-proficient ipNF95.11b C (**E, F**) and in the MPNST cell line ST-8814 (**G, H**) under the condition of MTAP knockdown. Cell lines were pretreated with 500 ng/mL doxycycline for 6 days to knock down MTAP in the +Dox group before inhibitor treatment. *p<0.05, **p<0.01, ***p<0.001, ****p<0.0001. **I**. Overall inhibitory effects of the two PRMT5 inhibitors in the ipNF95.11b C and ST-8814 cell lines with and without MTAP knockdown. The areas under the curves (AUCs) for the untreated control group and the treatment groups were compared in the presence or absence of MTAP knockdown, respectively. The initial cell counts measured on day 0 served as the baselines for the calculation. ****p<0.0001.

### PRMT5 overexpression promoted cell growth of MTAP-deficient benign neurofibroma cells

Our data suggest that MTAP loss leads to increased PRMT5 expression, likely as an adaptation in cells that lose MTAP and CDKN2A expression during malignant transformation ^6^. To test this hypothesis, we turned to neurofibroma cell lines with varying MTAP levels: MTAP-proficient ipNF95.11b C, MTAP-reduced ipn02.3 2λ and ipnNF95.11c (about 70-80% reduced compared to ipNF95.11b C) (Fig. 6A). First, we created stable cell lines overexpressing the human *PRMT5* gene using a doxycycline-inducible lentiviral system. This system enabled us to control PRMT5 expression (Fig. 6B-C) tightly. We then examined cell growth using the MTT assay. As expected, PRMT5 overexpression promoted cell growth of the MTAP-deficient ipn02.3 2λ and ipnNF95.11c but didn’t significantly affect the MTAP-proficient ipNF95.11b C (Fig. 6D-F). Consistently, PRMT5 enhanced the single-cell colony formation ability of the MTAP-deficient ipn02.3 2λ and ipnNF95.11c, but not the MTAP-proficient ipNF95.11b C(Fig. 6G-H). Thus, our results suggested that PRMT5 overexpression is a growth compensatory mechanism for MTAP loss at the molecular level.

**Figure 6.**
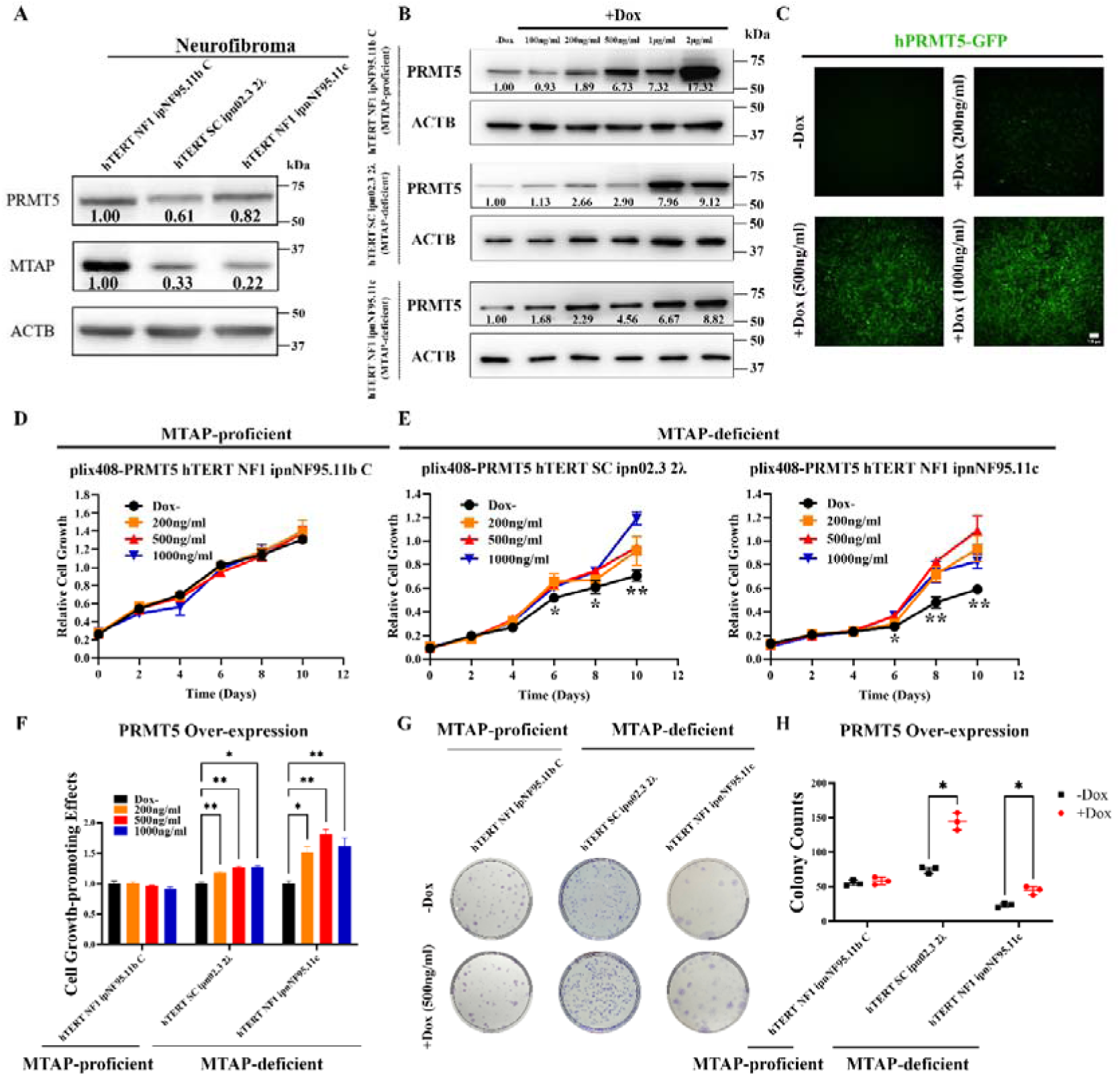
PRMT5 overexpression promoted cell growth of MTAP-deficient benign neurofibroma cells. **A**. Endogenous PRMT5 and MTAP in the 2 transformed immortalized neurofibroma cell lines that had relatively low MTAP ratios compared with the control cell line hTERT NF1 ipNF95.11b C. **B**. Validation of PRMT5 over-expression induced by a series of doxycycline concentration (100-2000ng/ml) in the 3 neurofibroma cell lines. **C.** The doxycycline simultaneously induced PRMT5 and GFP co-expression in the hTERT NF1 ipNF95.11b C cell line. **D**-**E**. Effects of the PRMT5 overexpression on the overall cell growth of neurofibroma cell lines by MTT assay. Cell counts for the control group (black: -Dox) and the PRMT5 overexpression group (red: +Dox) were evaluated every 2 days for a total of 10 days. OD595 values were measured for three replicates per condition. *p<0.05. **F**. Overall PRMT5 overexpression effects in the cell lines were evaluated by comparing the area under the curves (AUCs) of the -Dox control group and the +Dox treatment group. The initial cell counts measured on day 0 served as the baseline for the calculation. *p<0.05, ***p<0.001. ns. Nonsignificant. **G**-**H**. Effects of the PRMT5 overexpression on colony formation of the MTAP-deficient transformed neurofibroma cell lines. Doxycycline treatment was administered for 8 to 12 days. Colonies were visualized via crystal violet staining. The colonies with diameters> 0.2 μm were counted. Three replicates were performed for each group. *p<0.05, **p<0.01. ns. Nonsignificant.

### Slow cell growth of PRMT5 inhibition is mainly due to G2/M cell cycle arrest

The effects of PRMT5 knockdown on cell growth and colony formation could be due to a slowed cell cycle or increased cell death. Because PRMT5 has been reported to regulate the cell cycle ^60,61^, we first assessed cell cycle progression by flow cytometry through the 12^th^ day PRMT5 knockdown. An increased G2/M cell proportion was observed in the MTAP-deficient STS-26T and sNF96.2 after the 10^th^ day knockdown, but not in the MTAP-proficient ST-8814 and STS-26T, even on the 12^th^ day (Fig. 7A-B). In addition, increased G2/M cell cycle subpopulations were observed in PRMT5 chemical inhibitor JNJ-64619178 treatment assays, with consistent results in MTAP-deficient MPNST cells but not in MTAP-proficient MPNST cells (Fig. 7C-D). This increased G2/M was confirmed by the Fucci cell cycle assay, in which the increase in the GFP-labeled G2/M subpopulation is significant (Fig. 7E-F, green cells in the Fucci labeling). Next, we examined apoptosis with Annexin-V and senescence with β-galactosidase staining. We did not find an increase in cell death at the 12^th^ day after PRMT5 knockdown (Fig. S4). Thus, we reasoned that cell growth inhibition was mainly due to slow proliferation resulting from G2/M cell cycle arrest.

**Figure 7.**
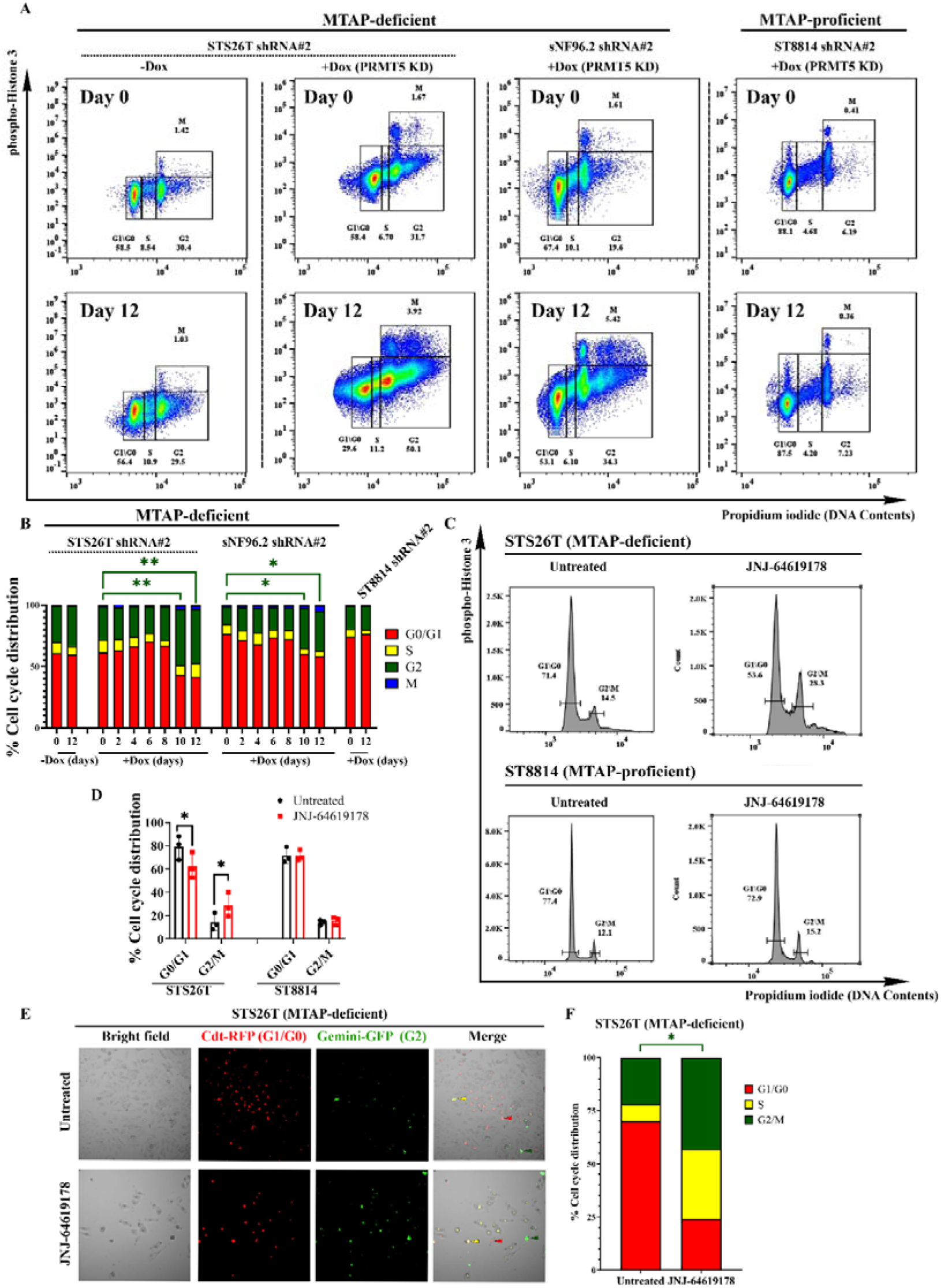
PRMT5 inhibition caused G2/M cell cycle arrest in the MTAP-deficient MPNST cell lines. **A**. Cell cycle analysis of -Dox and +Dox treatment for 12 days using flow cytometry. Cells were stained with propidium iodide (PI) and mitosis marker phospho-Histone H3 in the MTAP-deficient (STS-26T and sNF96.2) and MTAP-proficient (ST-8814) MPNST cell lines. G_1_/G_0_, G_2_, and M subpopulations were labeled with boxes. 500 ng/mL doxycycline was administered to induce shRNA#2-mediated PRMT5 knockdown. **B**. quantification of the subpopulations of the different cell cycle stages from day 0 to day 12. Three replicates were performed, and the averages were presented for each measurement. **C**. Cell cycle at the 3^rd^ day with and without 0.01μM JNJ-64619178 treatment by DNA content staining dye propidium iodide (PI) in the MTAP-deficient, STS-26T, and MTAP-proficient, ST-8814, MPNST cell lines. G1/G0 and G2/M subpopulations were labeled with the horizontal bars. **D**. Quantification of the subpopulation of the cell cycle in the STS-26T and ST-8814 with and without 0.01μM JNJ-64619178 treatment. Three replicates were performed for each measurement. *p<0.05. ns. Nonsignificant. **E**. Cell cycle analysis by the Fucci assay in the STS-26T cell line with and without PRMT5 inhibitor 0.01μM JNJ-64619178 treatment for 3 days. Cdt-RFP labeled G1 subpopulation, Gemini-GFP labeled G2/M subpopulation, and double positive cells represented the S stage subpopulation. **F**. Cell cycle stage quantification of Fucci assays. Five microscope view fields were randomly selected, and the average percentages of G1, S, and G2/M subpopulations were calculated and displayed in the bar graph.

### The DNA damage response likely causes G2/M cell cycle arrest after PRMT5 inhibition

The delayed cell growth beyond the 10^th^ day in response to PRMT5 inhibition led us to reason that the G2/M cell cycle arrest may be an indirect result of other cellular stresses, such as the DNA damage response, as reported in previous studies ^62^. To test this hypothesis, we first examined the DNA damage response by western blotting using antibodies against H2A.X and γ-H2A.X (phospho-H2A.X, Ser139), which are required for DNA repair ^63,64^. After PRMT5 knockdown, γ-H2A.X gradually accumulated, became evident on day 8, and was highest on day 12 in the MTAP-deficient STS-26T and sNF96.2, but not MTAP-proficient ST-8814 (Fig. 8A-F). To further confirm and visualize DNA damage, γ-H2AX staining by cellular immunofluorescence was performed on the 8^th^ day of PRMT5 knockdown. Indeed, there were more γ-H2AX-positive nuclear foci in the knockdown cells (Fig. 8G-H). As γ-H2A.X can be involved in many other processes, such as cell cycle arrest and cell death, we conducted Comet assays to quantify DNA damage in the PRMT5-inhibited STS-26T and sNF96.2 cells. DNA breaks were barely detectable before day 6 but became evident after day 8 following PRMT5 knockdown (Fig. 8I-L). Thus, PRMT5 inhibition led to the accumulation of DNA damage before G2/M cell cycle arrest, indicating that this accumulation is the primary cause of decreased cell growth.

**Figure 8.**
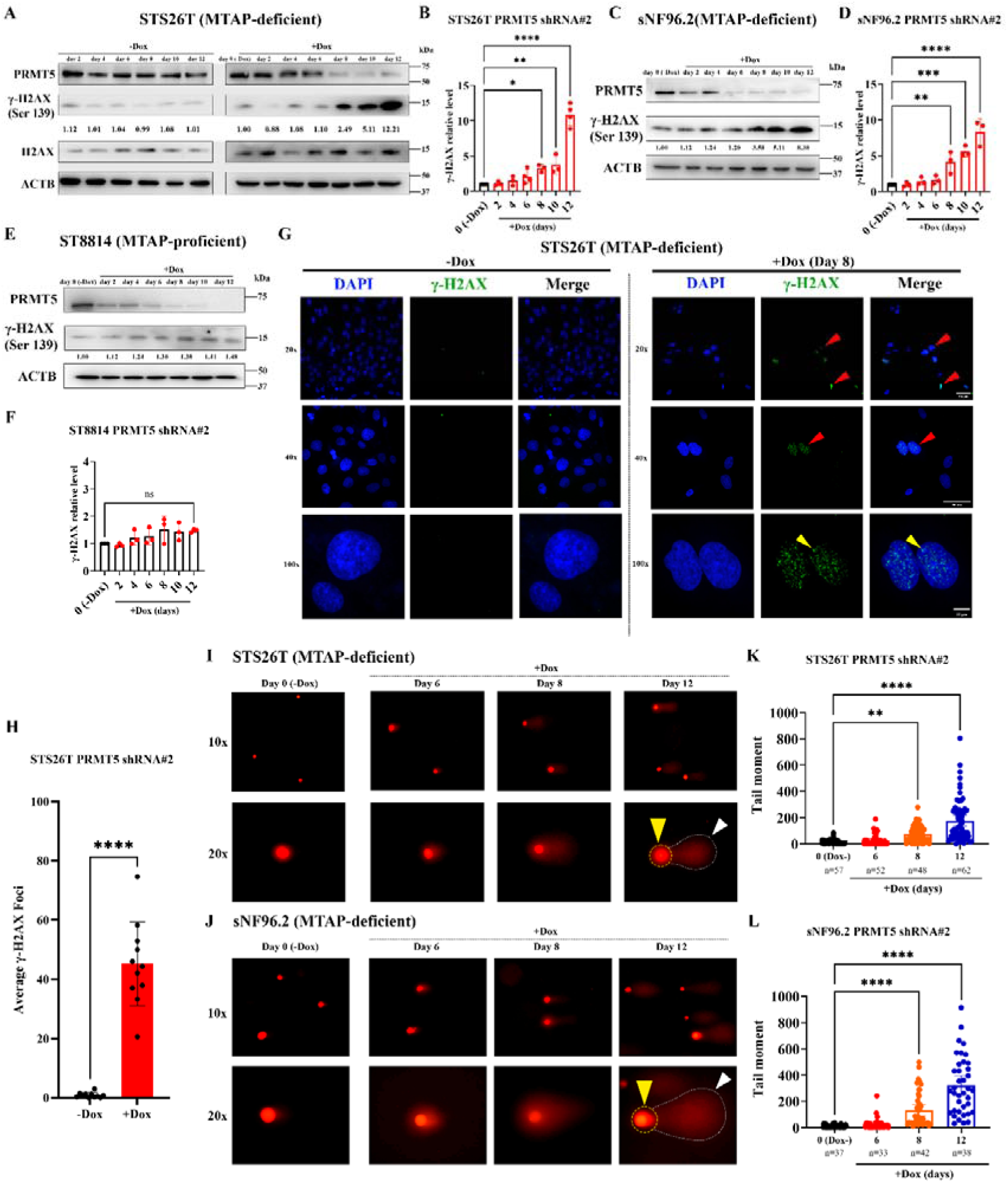
DNA damage accumulation preceded the G2/M cell cycle arrest after PRMT5 inhibition in the MTAP-deficient MPNST cell lines. **A**-**F**. DNA damage accumulation in the MPNST cell lines was measured by γ-H2AX (Ser139) using western blots. PRMT5 was knocked down by shRNA#2 in MTAP-deficient cell lines (STS-26T, sNF96.2) and the MTAP-proficient MPNST cell line (ST-8814). γ-H2AX levels were evaluated every 2 days from the start of Dox treatment (day 0) to day 12. The quantification of γ-H2AX levels at 0-12 days was compared to the doxycycline-untreated group. ACTB was used as the loading control. Three replicates were performed for statistical analysis. *p<0.05, **p<0.01, ***p<0.001, ****p<0.0001 **G-H**. Foci of γ-H2AX were detectable using immunofluorescence staining on the MTAP-deficient PRMT5 shRNA#2 STS-26T MPNST cell line. Cells were prepared by treatment with 500 ng/mL doxycycline for 8 days before staining and microscopy. Red arrowheads in F indicate nuclei staining, and the yellow arrowhead shows a γ-H2AX focus in a nucleus. Ten microscopic fields were randomly selected, and the average number of foci per nucleus in each field was calculated and displayed in the bar graph. ****p<0.0001. **I**-**L**. Measurement of accumulated DNA breaks by comet assay during the 6-12 days after PRMT5 knockdown. The tails, separated from the bright nucleus, indicate the extent of fragmented DNA, *i.e.*, DNA strand breaks. Quantification of DNA breaks is achieved by calculating the tail moment in the comet assays. Tail moments were calculated based on the tail brightness and the tail length. At least 25 comets were counted for each group. *p<0.05, ****p<0.0001. ns. Nonsignificant. **J**-**K**.

### Decreased RPA32 and activation of the phospho-CHK1 pathway following PRMT5 blockade

To identify the molecular mechanisms underlying DNA damage accumulation and cell cycle arrest after PRMT5 inhibition, we first examined expression levels of several proteins involved in DNA repair: RAD51 as a marker of homologous recombination, Ku70 and Ku80 for non-homologous end joining, and one of the common single-strand DNA repair markers, RAP32 (replication protein A subunit 32), in PRMT5 knockdown cells ^65–68^. Among these markers, RPA32 was consistently and significantly reduced by up to 70% in the two MTAP-deficient, not MTAP-proficient MPNST cells (Fig. 9A). Reduced RPA32 was observed at the 6^th^ day after PRMT5 knockdown in the STS-26T and sNF96.2, and this was earlier than DNA damage accumulation as well as cell cycle arrest (Fig. 9A-B, Fig. 8A-B). Using cellular immunofluorescence staining, we confirmed the decreased RPA32 in the PRMT5 knockdown STS-26T cell nucleus on the 6^th^ day (Fig. 9C). Thus, PRMT5 inhibition may cause DNA damage accumulation by reducing RPA32 in MTAP-deficient MPNST cells.

**Figure 9.**
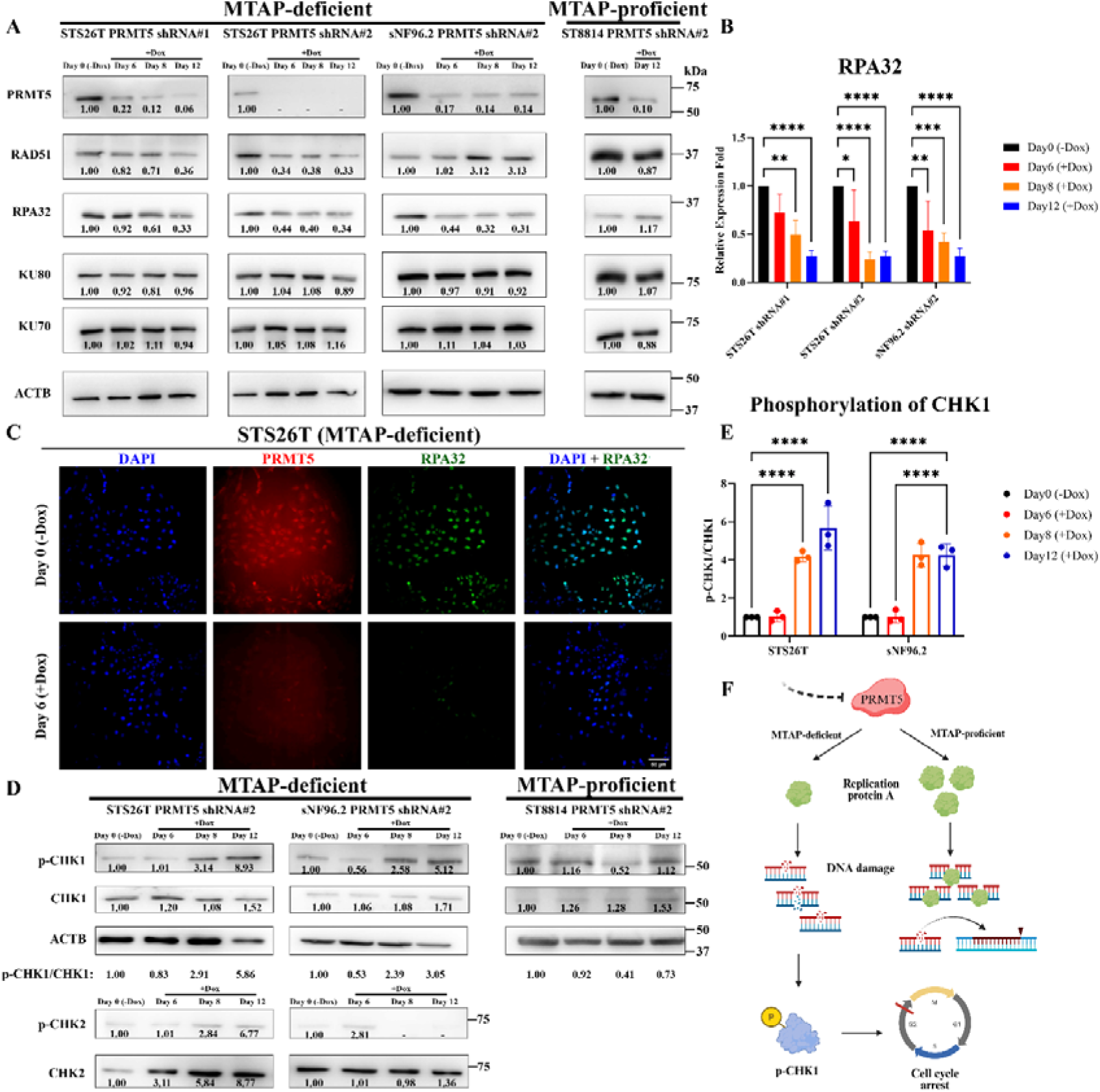
PRMT5 inhibition decreased RPA32 levels and activated the CHK1 signaling pathway in MTAP-deficient MPNST cells. **A**. Western blot analysis of DNA repair proteins in MPNST cell lines after PRMT5 knockdown. MTAP-deficient cell lines, STS-26T and sNF96.2, and the MTAP-proficient cell line, ST-8814, were analyzed from day 6 to day 12, after PRMT5 knockdown. The doxycycline-untreated group for each cell line served as the control. ACTB was used as the loading control. Three replicates were performed for statistical analysis. *p<0.05, **p<0.01, ***p<0.001. ****p<0.0001. **B**. Quantification of RPA32 protein levels in the MTAP-deficient MPNST cell lines STS-26T and sNF96.2 in the condition of PRMT5 knockdown from day 6 to day 12. **C**. Immunofluorescence staining of PRMT5 and RPA32 in the MTAP-deficient PRMT5 shRNA#2 STS-26T cell line at the 6^th^ day after PRMT5 knockdown. **D**. Western blot analysis of p-CHK1, CHK1, p-CHK2, and CHK2 from day 6 to day 12 after PRMT5 knockdown in the MTAP-deficient STS-26T and sNF96.2 and MTAP-proficient ST-8814 MPNST cell lines. **E**. Quantification of phosphorylation of CHK1 in the MTAP-deficient MPNST cell lines STS-26T and sNF96.2 cell lines from day 6 to day 12 after PRMT5 knockdown. Three replicates were performed for statistical analysis. ****p<0.0001. **F**. Illustration of PRMT5 mechanisms for cell growth inhibition in the MTAP-deficient MPNST cells after PRMT5 knockdown.

Reduced RPA32 may lead to accumulation of DNA breaks, which can result from impairing DNA replication or DNA damage repair, since the RPA complex is heavily involved in both ^68,69^. Given the evident G2/M cell cycle arrest, we then evaluated the CHK1 and CHK2 pathways for single- and double-strand breaks, respectively. CHK1 signaling was activated as determined by phosphorylation at Ser345 on the 8^th^ day after PRMT5 knockdown in MTAP-deficient, but not MTAP-proficient MPNST cells.

This coincided with the accumulation of DNA damage as measured by γ-H2AX (Fig. 8A-B, Fig. 9D-E). Thus, the increase in replication stress due to RPA deficiency led to activation of the DNA damage response via CHK1, with a resulting cell cycle arrest in the MTAP-deficient MPNST cells by PRMT5 knockdown (Fig. 9F). The absence of CHK2 phosphorylation indicates that DNA replication, instead of double-stranded DNA repair, is the primary cause of DNA damage accumulation.

### A combination of PRMT5 blockade and DNA damage-inducing chemotherapy drugs resulted in synergistic effects exclusively in MTAP-deficient MPNST cells

Although PRMT5 inhibition resulted in DNA damage accumulation during DNA replication and G2/M cell cycle arrest, apoptosis assays did not indicate cancer cell killing. To assess whether additional DNA damage in these cells would enhance specific killing efficiency, we selected three common chemotherapy drugs: doxorubicin, gemcitabine, and 5-FU (5-fluorouracil). Doxorubicin is the first-line chemotherapy drug for MPNSTs. Both doxorubicin and gemcitabine are DNA-damaging drugs commonly used in chemotherapy regimens, though with limited response and efficacy ^70,71^. The 5-FU served as a control as it primarily interrupts DNA synthesis and induces DNA replication stress, which is also influenced by PRMT5 inhibition. Then, we performed IC_50_ assays with MTAP-deficient and MTAP-proficient MPNST cell lines with or without induced PRMT5 knockdown by shRNA (Fig. 10A) and Cas13d (Fig. 10B). Indeed, PRMT5 inhibition largely sensitized MTAP-deficient MPNST cells to the DNA damage-inducing drugs, doxorubicin and gemcitabine, but not the MTAP-proficient MPNST cell line ST-8814 (Fig. 10A-B). An IC_50_ assay indicated that PRMT5 knockdown rendered MTAP-deficient MPNST cells 6- to 400-fold more sensitive to doxorubicin and gemcitabine (Fig. 10A-C). In contrast, PRMT5 inhibition didn’t have such a synergistic effect with 5-FU, which targets thymidylate synthase and inhibits DNA synthesis, rather than directly inducing DNA damage (Fig. 10A-C).

**Figure 10.**
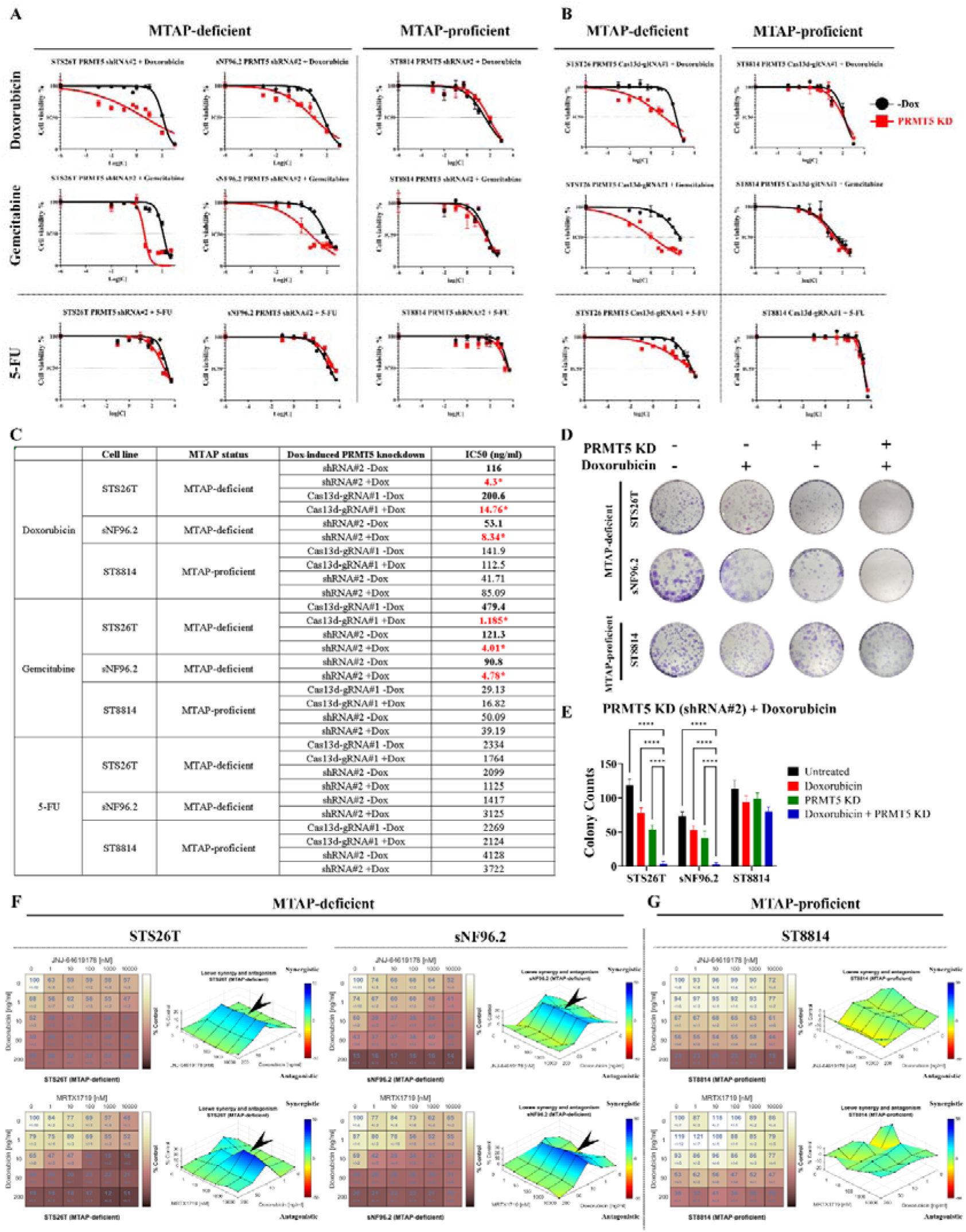
PRMT5 inhibition synergized with the DNA damage-inducing chemotherapy agents. **A**. IC_50_ analysis of the three chemotherapy agents in PRMT5 knockdown cells by shRNA. Drug concentration-response curves were generated based on the 10 observation points in the two MTAP-deficient cell lines, sNF96.2 and STS-26T, and an MTAP-proficient cell line, ST-8814, after PRMT5 inhibition by shRNA. Each observation point, consisting of three replicates, represented the normalized cell viability at a single concentration of doxorubicin, gemcitabine, or 5-FU relative to the corresponding untreated groups. A log (Concentration)-response model with variable slopes and four parameters was used. The cell lines were untreated (-Dox) or pretreated with 500 ng/mL Dox for 5 days, followed by chemotherapy drug treatment for 5 days. **B**. Validation of IC_50_ analysis of the three chemotherapy agents in PRMT5 knockdown cells by Cas13d. STS-26T and ST-8814 were analyzed. **C**. The estimated IC_50_ values of the three cell lines to doxorubicin, gemcitabine, and 5-FU are based on the drug concentration response curves. **D**-**E**. Combinational effect on the colony formation between PRMT5 knockdown and doxorubicin (10ng/mL). The Dox pre-treatment was administered for 5 days before doxorubicin treatment. Colonies were visualized via crystal violet staining. The colonies with diameters> 0.2 μm were counted. Three replicates were performed for each group. ***p<0.001. **F**-**G**. The synergistic effect index of the combination treatment of the four PRMT5 inhibitors and doxorubicin. The Loewe synergy and antagonism model was used to evaluate the combination treatment effects. The blue color indicates synergistic effects of the combination treatments. The index standard bar shows the highest synergy (blue), no interaction (green), and the highest antagonism (red).

IC_50_ assay indicated significantly lower effective concentrations of both doxorubicin and gemcitabine in MTAP-deficient MPNST cells under PRMT5 knockdown. Thus, we selected 10 ng/mL of these chemotherapy drugs and combined them with PRMT5 knockdown for the colony formation assay. We confirmed that the positive response to the combination treatment was observed in STS-26T and sNF96.2, but not in ST-8814 (Fig. 10D-E). Moreover, 10ng/mL doxorubicin combined with shRNA-mediated PRMT5 knockdown was able to induce cellular apoptosis, but neither doxorubicin nor PRMT5 knockdown alone did so in the MTAP-deficient MPNST cell STS-26T (Fig. S5). To further test the therapeutic potential of this synergy concept, we combined two PRMT5 chemical inhibitors, JNJ-64619178 and MRTX1719, with doxorubicin. Consistently, both PRMT5 inhibitors synergized with doxorubicin at 10 ng/mL on the MTAP-deficient MPNST cells, but not MTAP-proficient MPNST cells (Fig.10F-G). Thus, PRMT5 inhibition synergized with DNA damage-inducing drugs, potentially enhancing their therapeutic effects, as evidenced by increased apoptosis.

## DISCUSSION

MPNSTs are genetically heterogeneous tumors with poor prognosis due to frequent resistance to chemotherapy and radiotherapy, and a lack of effective targeted therapy. Here, we confirmed that frequent MTAP loss is associated with high PRMT5 expression in human MPNSTs. In addition, genetic and pharmacological inhibition of PRMT5 in MTAP-deficient MPNST cells disrupted DNA damage homeostasis, induced cell cycle arrest, and sensitized cells to DNA damage–inducing agents. These findings suggest that PRMT5 inhibition may be a new therapeutic target, with the potential for synergy in combination with DNA-damaging agents and potentially even more effective in patients with low or deleted MTAP.

With immunohistochemistry, we found a clear relationship between MTAP loss and PRMT5 overexpression not only in MPNSTs but also in NFs, despite the notorious histological heterogeneity. Compared with normal nerves, MTAP is frequently lost or downregulated in pathological conditions (NFs, ANNUBPs, and MPNSTs). Low-grade MPNSTs exhibited some MTAP expression, likely due to stochastic effects given the small sample size. Future studies are needed to clarify MTAP expression in these samples further. MTAP loss in neurofibroma is consistent with the loss of *CDKN2A* on chromosome 9p during the transition from benign to malignant ^6,7^. Since MTAP loss and PRMT5 overexpression are common in human MPNSTs, clinical evaluation of tumor MTAP loss compared with adjacent normal nerve tissue could be valuable for stratifying patients who might be sensitive to PRMT5 inhibition.

Although human cancer cell lines are widely used to investigate the biology of the corresponding cancers, our analyses revealed discrepancies between cell lines and cancer tissues. Among 12 currently available MPNST cell lines, we found that *CDKN2A* (P16) was lost in most (10 of 12), MTAP deficiency occurred in approximately half (6 of 12), and none showed strong PRMT5 expression, even in MTAP-deficient cell lines, unlike human MPNSTs. This difference could be attributed to the use of enriched cell culture media or selective pressure during the establishment of the cell line. Nonetheless, the MTAP-deficient and MTAP-proficient cell lines responded distinctly to genetic and pharmacologic inhibition of PRMT5. Following blockade of PRMT5, MTAP-deficient cell growth was significantly slowed down compared to the MTAP-proficient cell lines. Consistently, the MTAP-proficient cell lines became sensitive to PRMT5 inhibitors upon MTAP knockdown. Cell growth decreased relatively late (beyond 8 days), and cell death was not evident, indicating that PRMT5 inhibition alone may not be sufficient to kill the cancer cells, and that the cells may undergo a new physiological adaptation through unknown mechanisms.

MTAP-PRMT5 is a well-known example of classical collateral lethality ^22,23,72^. The functional role of PRMT5 in MTAP-loss cancer cells remains a topic of debate. It has been proposed that reduced PRMT5 activity in MTAP-lost cancer cells renders them more vulnerable to additional inhibition, as the initial hit was imposed by endogenous MTA inhibition ^72,73^. However, this view cannot explain why PRMT5 is frequently overexpressed across many cancer types. In our study, we found a negative correlation between functional MTAP and PRMT5 in human MPNSTs as well as prompt compensatory upregulation of PRMT5 in response to MTAP knockdown in the MTAP-proficient cells. Moreover, PRMT5 overexpression promoted growth in the NF cell lines. Thus, we suggest that PRMT5 expression is regulated by a feedback mechanism to compensate for MTAP loss and support cell growth, rather than through a gain-of-function mutation. It is also worth noting that NF cell lines upregulated PRMT5 following MTAP knockdown, and they became sensitive to PRMT5 inhibitors. Thus, the PRMT5 inhibition strategy can also be applied to NF patients (with MTAP deletion) to prevent MPNST transition, given the prevalence of NF in the human population and the key role of *CDKN2A-MTAP* loss.

PRMT5 has already been shown to regulate DNA damage repair. For example, PRMT5 was reported to promote homologous recombination by methylating RUVBL1, a coactivator of TIP60, which is involved in the early DNA damage response to double-strand breaks ^74^. In our experiments, neither RAD51 (for homologous recombination) nor Ku70 and Ku80 (for non-homologous end-joining) were altered. Instead, we found that RAP32 decreased and CHK1 was activated, suggesting that PRMT5 inhibition causes DNA replication stress in MTAP-deficient cells. Coincidentally, a recent report also found PRMT5-dependent depletion of RPA32 in pancreatic cancer cells ^75^. These data indicate that PRMT5 plays a critical but poorly defined role in regulating RPA32. Moreover, DNA replication stress could be the leading cause of DNA break accumulation and cell cycle arrest following PRMT5 inhibition in MTAP-deficient cells. The mechanistic link between PRMT5 and RPA32 remains to be explored. It is likely that PRMT5 directly methylates RPA32, which is one of the three subunits of the replication protein A (RPA) complex. Indeed, a recent study reported mono-methylation of RPA70 (RPA1) by mass spectrometry *in vitro*, although it did not identify the responsible enzymes or functionally characterize the roles of the methylation modification within the RPA complex ^76^. With AlphaFold2, we found that RPA32 contained two intrinsically disordered regions (IDRs), and three potential arginine dimethylation sites were predicted within these IDRs (Fig. S5A-B). Future studies are needed to clarify how PRMT5 regulates RPA32.

The effect of PRMT5 inhibition depends on MTAP abundance in MPNST cells. Neither genetic nor chemical inhibition of PRMT5 killed the MTAP-deficient tumor cells. However, the killing efficiency was drastically increased by combined treatment with damage–inducing agents (doxorubicin and gemcitabine), but not with the thymidylate synthase (TS) inhibitor 5-FU, revealing that PRMT5 inhibition and damage–inducing agents synergized. Interestingly, a recent study showed an additive effect of PRMT5 inhibitors (TNG462 or TNG908) with chemotherapy drugs (doxorubicin or trabectedin) in both MTAP-null and MTAP-WT MPNST cell lines ^73^. In contrast, our results showed that PRMT5 inhibition specifically sensitized cancer cells, not cells with MTAP, to damage–inducing agents, leading to greater cell death. The discrepancy may be due to differences in the cell line’s genetic background and the chemical inhibitors used. Nonetheless, the study also supported the potential of PRMT5 inhibition in MPNSTs, using MPNST PDX lines and the mouse xenograft model ^73^.

Since several PRMT5 inhibitors have already entered clinical trials, the PRMT5 inhibition strategy has strong translational potential for MPNSTs. We expect that PRMT5 inhibition will enter clinical practice for MPNSTs and other MTAP-deficient cancers, likely in combination with chemotherapy, radiotherapy, or other targeted inhibitors. Indeed, this type of PRMT5 synergy has been reported in pancreatic cancer ^75,77^. PRMT5 inhibitors were found to be useful in CDK4/6 inhibitor-resistant ER+/RB-deficient breast cancer ^78^ and to improve response to PARP inhibitors ^79^. Additionally, paclitaxel was discovered to overcome resistance to PRMT5 inhibition, demonstrating the applicability of a rationale combination therapy that includes PRMT5 inhibitors ^77^. As the standard of care for MPNST is doxorubicin/ifosfamide, and PRMT5 inhibitors synergize with doxorubicin, these data pave the way for future clinical applications.

## ACKNOWLEDGEMENT

This work was primarily supported by the CDMRP Neurofibromatosis Research Program under Award No. W81XWH-22-1-0265. The lab and part of the work were also supported by the National Institute of General Medical Sciences of the National Institutes of Health (R35GM124913), the Heyward Foundation, and Purdue University College of Veterinary Medicine. The content is solely the responsibility of the authors and does not necessarily represent the official views of the funding agents. We also thank the Neurofibromatosis Therapeutic Acceleration Program (NTAP) for NF1 cell lines and Dr. Christine A. Pratilas’ lab at Johns Hopkins University for providing some MPNST cell lines.

## AUTHOR CONTRIBUTIONS

DW and GZ analyzed the data and wrote the first draft of the manuscript. DW performed experiments. MLF and SG contributed to the establishment of some MPNST cell lines. AS contributed to the collection of paraffin-embedded samples and related information. DW, CH, SC, and GZ conceived the concept and designed experiments. CH, SC, and GZ managed the project and acquired research funding. All authors reviewed and edited the final manuscript.

