## Supplementary file for "PRMT5 is Frequently Upregulated and a Potential Therapeutic Target in MTAP-deficient Malignant Peripheral Nerve Sheath Tumors"

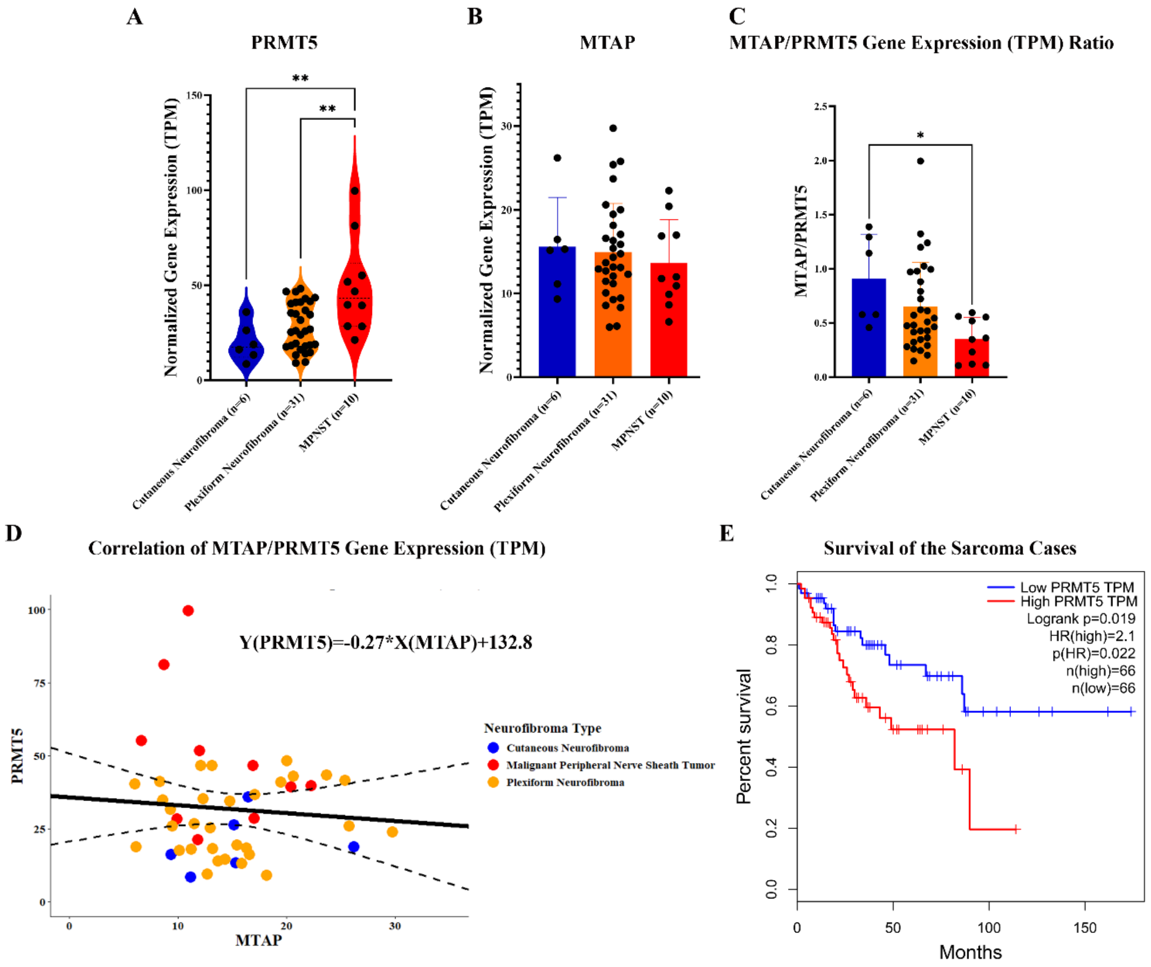


**Figure S1. PRMT5 and MTAP expression changes in MPNST and other types of cancer. A-B.** PRMT5 and MTAP mRNA levels in the cutaneous neurofibroma, plexiform neurofibroma, and MPNST. The whole-exome sequencing data contributed by Johns Hopkins University were obtained from cBioportal (https://www.cbioportal.org/). Transcripts per million (TPM) was used to quantify and compare the mRNA levels. **C.** Ratios of MTAP and PRMT5 mRNA levels (TPM) in the cutaneous neurofibroma, plexiform neurofibroma, and MPNST. **D.** General linear correlation between MTAP and PRMT5 mRNA levels (TPM) in the cutaneous neurofibroma, plexiform neurofibroma, and MPNST. **E.** Kaplan-Meier survival analysis of the sarcoma patients stratified with PRMT5 mRNA levels (TPM). The highest and lowest quartiles of PRMT5 mRNA levels (TPM) among all sarcoma patients defined the high- and low-PRMT5 expression groups, respectively.

**PRMT5 is Frequently Upregulated and a Potential Therapeutic Target in MTAP-deficient Malignant Peripheral Nerve Sheath Tumors**

The data were obtained from the TCGA dataset and analyzed using GEPIA. HR: hazard ratio.

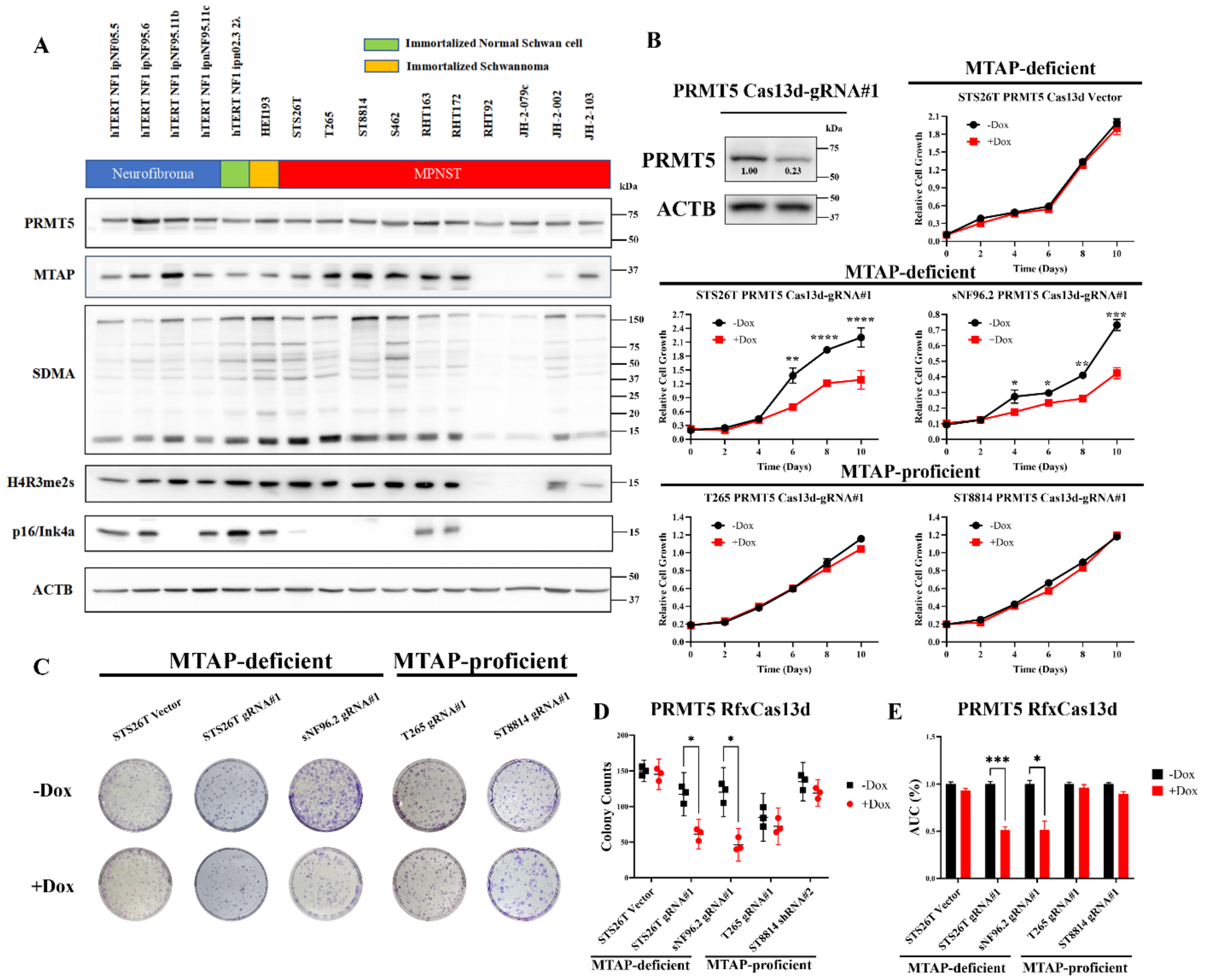


**Figure S2. Cas13d-mediated PRMT5 knockdown inhibited MTAP-deficient MPNST cell growth. A.** Endogenous levels of PRMT5, MTAP, symmetrical di-methylation arginine (SDMA), H4R3me2s, p16 (CDKN2A), and ACTB in the immortalized neurofibroma/schwannoma and MPNST cell lines. **B.** PRMT5 knockdown mediated by RfxCas13d-gRNA#1 in the MPNST cell lines. Western blot showed PRMT5 expression levels in the STS26T cell line with or without 3-day doxycycline (Dox, 500 ng/mL) treatment. The gRNA efficiency was quantified by comparing PRMT5 expression levels in the Dox+ and Dox- groups. The effects of the PRMT5 Cas13d-gRNA#1 or the vector on the overall cell growth of the MPNST cell lines were evaluated by MTT assay. Cell counts of the control group (black: Dox-) and the PRMT5 knockdown group (red: +Dox)

were evaluated every 2 days for a total of 10 days. $OD_{595}$ values were measured from three replicates per condition. *p<0.05, **p<0.01, ****p<0.0001. **C.** PRMT5 knockdown effects by gRNA#1 or the empty vector control of the MPNST cell lines were assessed by colony formation assays. Dox treatment was administered for 6-12 days. The colony was visualized by crystal violet staining. The colony with a diameter> 0.2 μm was counted. Three replicates were performed for each group. *p<0.05, **p<0.01, ***p<0.001. **D**. Quantification and statistical analysis of the colony formation assays. **E**. Quantification of the inhibitory effects of PRMT5 RfxCas13d-gRNA#1 or vector in the MTAP-deficient and -proficient MPNST cell lines by using AUCs from growth curves in panel B.

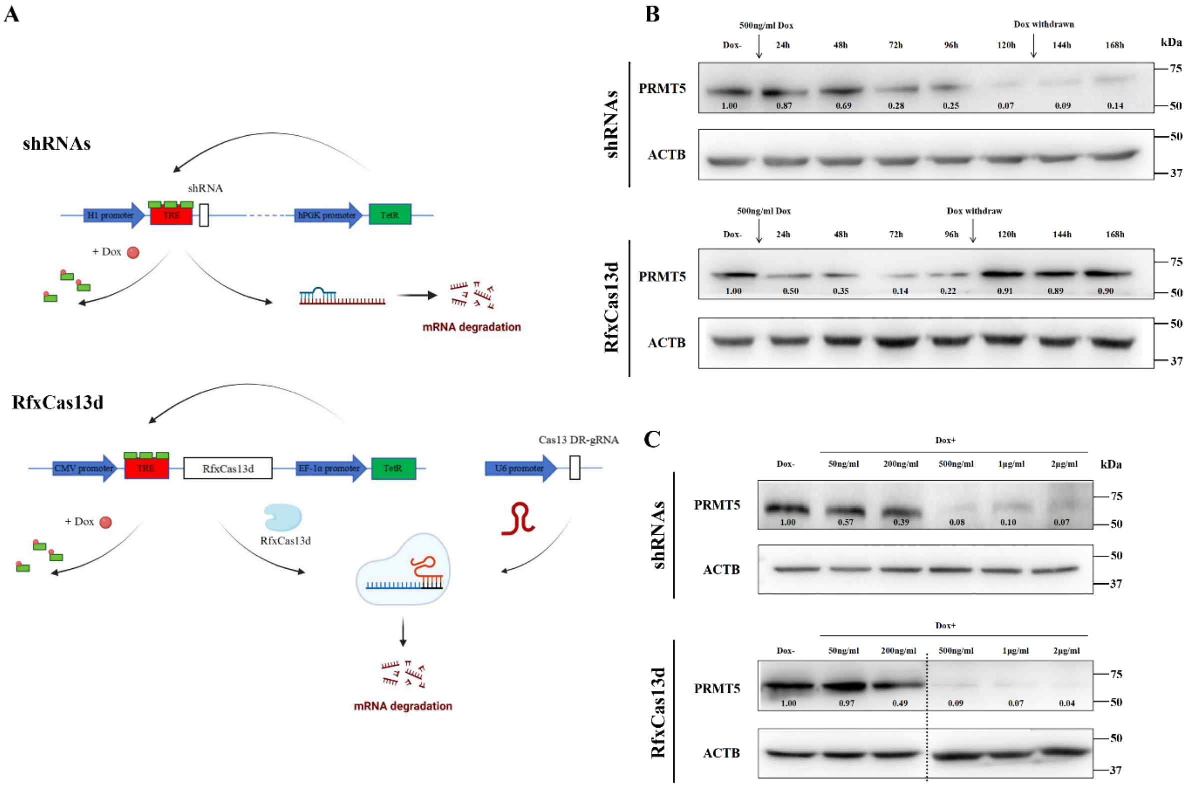


**Figure S3. PRMT5 knockdown by shRNA and RfxCas13d. A.** The illustration of inducible design of PRMT5 knockdown mediated by shRNA and RfxCas13d. **B.** The PRMT5 knockdown efficiency by PRMT5 shRNA#2 and PRMT5 RfxCas13d-gRNA#1 in the STS26T cell line during the time course of 7 days. **C.** The PRMT5 knockdown efficiency by PRMT5 shRNA#2 and PRMT5 RfxCas13d-gRNA#1 in the STS26T cell line at the different doxycycline concentrations for 5 days.

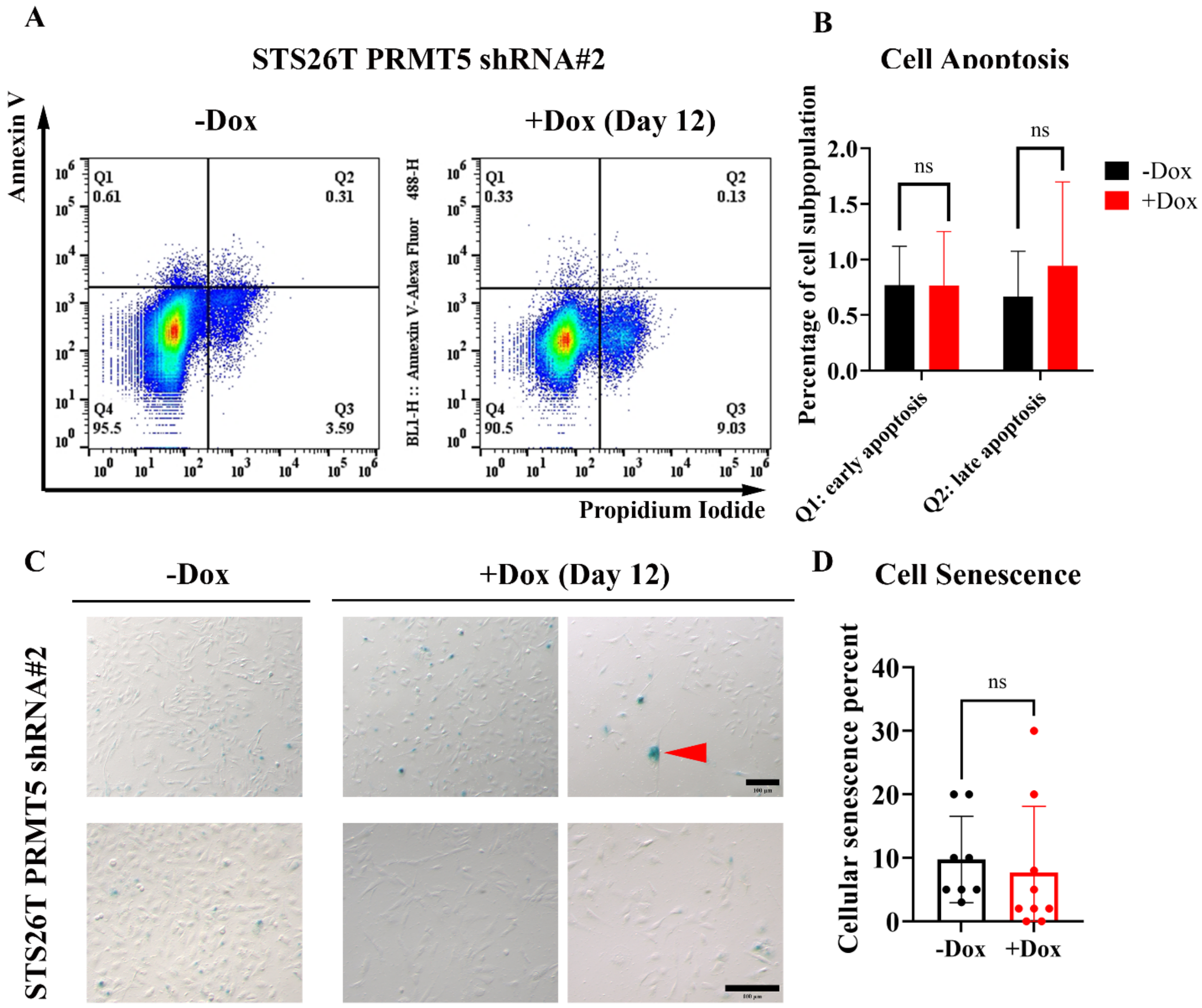


**Figure S4. PRMT5 knockdown didn't induce cellular senescence and apoptosis in the MTAP-deficient MPNST cells. A-B.** Flow cytometry with Annexin V and propidium iodide staining to detect cellular apoptosis in the PRMT5 shRNA#2 MTAP-deficient STS26T MPNST cell line. 500ng/mL doxycycline was administered for 12 days. Three replicates were performed for each group. Q1 (early apoptosis) and Q2 (late apoptosis) sub-populations were quantified for statistical analysis. **C-D.** Cellular senescence measured by β-galactosidase staining on the PRMT5 shRNA#2 MTAP-deficient STS26T MPNST cell line. 500 ng/mL doxycycline was administered for 12 days. A minimum of five randomly selected fields per well were imaged. Each group has six wells. The red arrow ahead indicates a positive-staining cell. The average percentages of blue-positive cells were calculated and presented in a bar graph for statistical analysis.

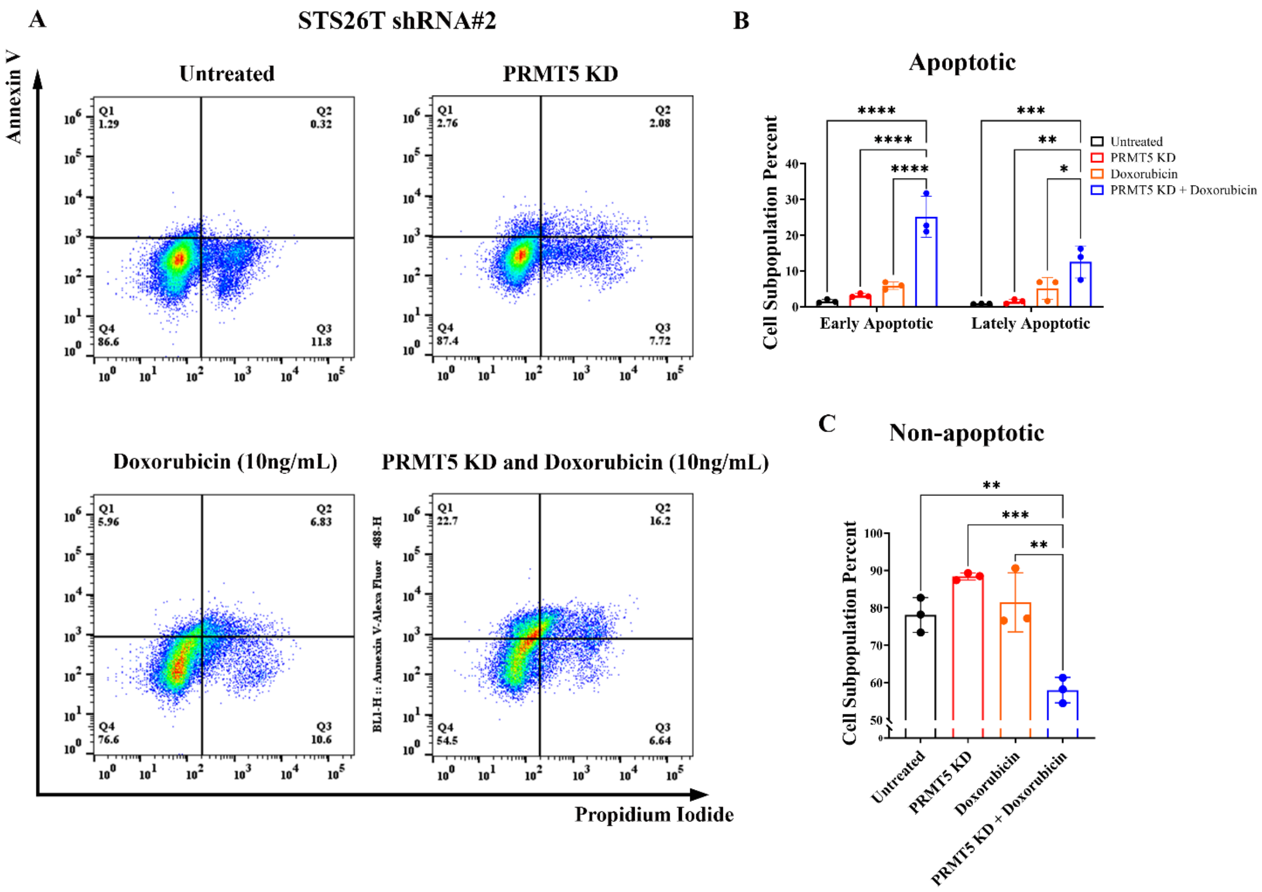


**Figure S5. Combination treatment of PRMT5 inhibition and doxorubicin induced cell apoptosis in the MTAP-deficient MPNST cells. A.** Flow cytometry with Annexin V and propidium iodide staining to detect cellular apoptosis in the PRMT5 shRNA#2 MTAP-deficient STS26T

MPNST cell line. 500ng/mL doxycycline was administered for 5 days followed by 10ng/mL doxorubicin for 2 days. Three replicates were performed for each group. **B-C.** Quantification and comparison of apoptotic (Q1 and Q2) and non-apoptotic (Q4) cell sub-populations.

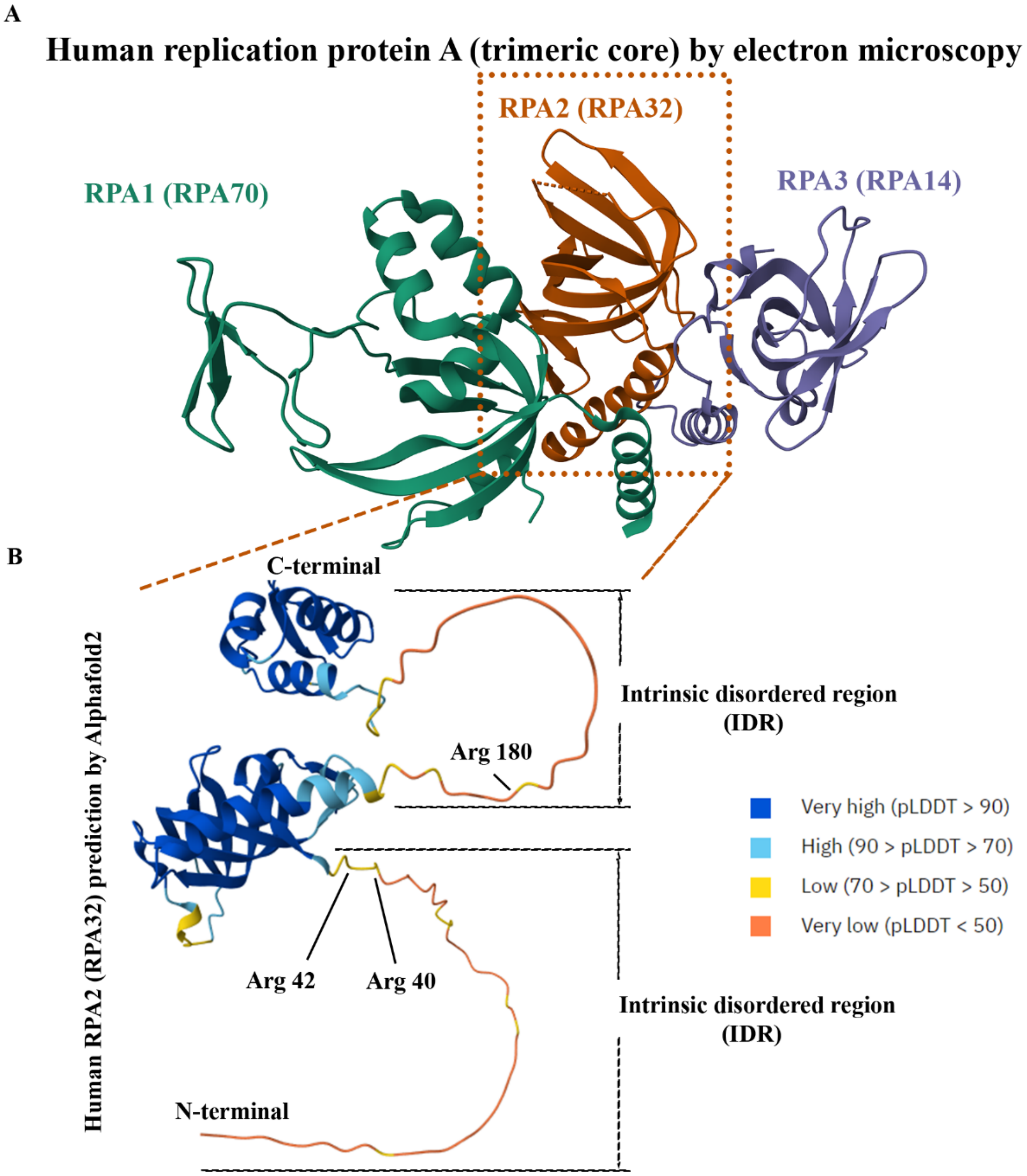


**Figure S6. Potential symmetrical di-methylation arginine sites in the RPA32. A.** Human RPA complex structure by electron microscopy. The protein tertiary structure data were from the Protein Data Bank (https://doi.org/10.2210/pdb8rk2/pdb). Three subunits of the RPA complex, RPA70, RPA32, and RPA14, were highlighted in different colors. **B.** Human RPA32 predicted structure with potentially symmetrically di-methylated arginines by Alphafold2. The structure prediction (AF-A0A6A5SIS8-F1-v6) was obtained from the AlphaFold Protein Structure Database (https://alphafold.ebi.ac.uk/). Intrinsic disordered regions (IDRs) were labeled. The top three PRMT5-potentially dimethylated arginines, as identified by Alphafold2, were highlighted (Arg 40, Arg 42, and Arg 180).

**Supplement Table 1. MTAP Loss in the Atypical Neurofibroma and MPNST with the Clinicopathological Features**

| aNF and MPNST cases | | MTAP completely lost<br>23 | MTAP partially lost<br>10 | MTAP proficient<br>5 | Chi-square or Fisher's test p-value | N<br>38 |
|---|---|---|---|---|---|---|
| Clinical features | | | | | | |
| Gender | Male | 10 (66.7%) | 3 (20%) | 2 (13.3%) | 0.8021 | 15 |
| | Female | 12 (54.5%) | 7 (31.8%) | 3 (13.6%) | | 22 |
| Grade | aNF | 4 (80%) | 1 (20%) | 0 (0%) | 0.0746* | 5 |
| | Low-grade MPNST | 3 (50%) | 0 (0%) | 3 (50%) | | 6 |
| | High-grade MPNST | 16 (59.3%) | 9 (33.3%) | 2 (7.4%) | | 27 |
| Primary tumor site | Extremity | 6 (60%) | 2 (20%) | 2 (20%) | 0.8665 | 10 |
| | Trunk | 10 (58.8%) | 5 (29.4%) | 2 (11.8%) | | 17 |
| Metastasis | Metastasis occurred | 6 (60%) | 3 (30%) | 1 (10%) | 0.9979 | 10 |
| | No metastasis | 8 (53.3%) | 4 (26.7%) | 3 (20%) | | 15 |
| Recurrence | Recurred | 5 (45.5%) | 4 (36.4%) | 2 (18.2%) | 0.8598 | 11 |
| | Not recurred | 8 (61.5%) | 3 (23.1%) | 2 (15.4%) | | 13 |
| NF1 | *NF1* mutation | 12 (52.2%) | 7 (30.4%) | 4 (17.4%) | 0.8352 | 23 |
| | No *NF1* mutation | 4 (66.7%) | 2 (33.3%) | 0 (0%) | | 6 |
| Arising from pNF or NF | NF | 6 (50%) | 4 (33.3%) | 2 (16.7%) | 0.9894 | 12 |
| | No NF | 6 (60%) | 3 (30%) | 1 (10%) | | 10 |
| Histology | Spindle | 17 (56.7%) | 9 (30%) | 4 (13.3%) | N/A | 30 |
| | Epithelial | 1 (100%) | 0 (0%) | 0 (0%) | | 1 |
| Differentiation | Conventional | 12 (54.5%) | 6 (27.3%) | 4 (18.2%) | N/A | 22 |
| | Triton | 1 (50%) | 1 (50%) | 0 (0%) | | 2 |

*$p<0.1$

**Table S1. MTAP loss in atypical neurofibroma and MPNST, with clinicopathological features.**

### PRMT5 is Frequently Upregulated and a Potential Therapeutic Target in MTAP-deficient Malignant Peripheral Nerve Sheath Tumors

**Table 2. Univariate Analysis of PRMT5 Expression in the aNF and MPNST Patients**

| | | PRMT5 Expression (Mean +SD) | Student t-test/Mann-Whitney U or One-way Anova p-value | N |
|---|---|---|---|---|
| aNF and MPNST cases | | | | 38 |
| Clinical features | | | | |
| Gender | Male | 199 ± 64.54 | 0.6766 | 15 |
| | Female | 188.1 ± 76.46 | | 21 |
| Grade | aNF | 225 ± 42.43 | 0.4105 | 5 |
| | Low-grade MPNST | 185 ± 69.5 | | 6 |
| | High-grade MPNST | 190 ± 75.21 | | 26 |
| Primary tumor site | Extremity | 190 ± 93.67 | 0.4002 | 9 |
| | Trunk | 180.88 ± 68.1 | | 17 |
| Metastasis | Metastasis occurred | 198 ± 89.67 | 0.8278 | 10 |
| | No metastasis | 165 ± 69.48 | | 14 |
| Recurrence | Recurred | 191 ± 69.83 | 0.7219 | 10 |
| | Not recurred | 171.54 ± 86.2 | | 13 |
| NF1 | *NF1* mutation | 188.86 ± 60.35 | 0.706 | 22 |
| | No *NF1* mutation | 160 ± 119.16 | | 6 |
| Arising from pNF or NF | NF | 177.27 ± 66.8 | 0.3774 | 11 |
| | No NF | 189 ± 97.8 | | 10 |
| Histology | Spindle | 185.5 ± 73.16 | N/A | 30 |
| | Epithelial | N/A | | 0 |
| Differentiation | Conventional | 179.05 ± 76.81 | 0.4259 | 21 |
| | Triton | 160 ± 113.14 | | 2 |

**Table S2. Univariate analysis of PRMT5 expression in the atypical neurofibroma and MPNST patients.**

**Table 3. Univariate Analysis of survival time from diagnosis to death in the aNF and MPNST Patients**

| | | Survival time from Diagnosis to Death (Mean +SD, Days) | Student t-test/Mann-Whitney U test or One-way Anova p-value | N |
|---|---|---|---|---|
| aNF and MPNST cases | | | | 38 |
| Clinical features | | | | |
| Gender | Male | 459.25 ± 353.79 | 0.4056 | 4 |
| | Female | 716.2 ± 756.24 | | 10 |
| Grade | aNF | N/A | N/A | 0 |
| | Low-grade MPNST | 350 ± NA | | 1 |
| | High-grade MPNST | 455.33 ± 492.99 | | 9 |
| Primary tumor site | Extremity | 425.5 ± 89.5 | 0.7037 | 4 |
| | Trunk | 530.5 ± 740.22 | | 8 |
| Metastasis | Metastasis occurred | 238 ± 185.71 | 0.0791* | 6 |
| | No metastasis | 949.5 ± 790.17 | | 6 |
| Recurrence | Recurred | 282 ± 180.77 | 0.0455* | 6 |
| | Not recurred | 1123.83 ± 781.37 | | 8 |
| NF1 | *NF1* mutation | 584.2 ± 709.82 | 0.9039 | 10 |
| | No *NF1* mutation | 543.33 ± 406.63 | | 3 |
| Arising from pNF or NF | NF | 624.25 ± 797.67 | 0.6956 | 8 |
| | No NF | 490.75 ± 348.27 | | 4 |
| Histology | Spindle | 574.77 ± 637 | N/A | 13 |
| | Epithelial | N/A | | 0 |
| Differentiation | Conventional | 463.91 ± 435.36 | N/A | 11 |
| | Triton | 161 ± NA | | 1 |

*p<0.1

**Table S3. Univariate analysis of survival time from diagnosis to death in the atypical and MPNST patients.**

#### PRMT5 is Frequently Upregulated and a Potential Therapeutic Target in MTAP-deficient Malignant Peripheral Nerve Sheath Tumors

**Supplement Table 4. The Information of the Cell Line Collection**

| Cell Line Name | Cell Line Category | Germline neurofibromatosis status | Somatic mutations | Cell Morphology | References |
|---|---|---|---|---|---|
| hTERT NF1 ipNF05.5 | Plexiform neurofibroma, transformed cell line | *NF1:* c.3456_3457 | *NF1* LOH | Fibroblast/spindle | (Li et al., 2016) |
| hTERT NF1 ipNF95.6 | Plexiform neurofibroma, transformed cell line | *NF1:* R816X | *NF1:* R2237X | Fibroblast/spindle | (Li et al., 2016) |
| hTERT NF1 ipNF95.11b | Plexiform neurofibroma, transformed cell line | *NF1:* c.1756-delACTA | | Fibroblast/spindle | (Li et al., 2016) |
| hTERT NF1 ipnNF95.11c | Plexiform neurofibroma, transformed cell line, NF1 donor | *NF1:* c.1756-delACTA | | Fibroblast/spindle | (Li et al., 2016) |
| hTERT NF1 ipn02.3 2λ | Human Schwann cell, transformed cell line, non-NF1 donor | No NF1/*NF1* wild type | | Fibroblast/spindle | (Li et al., 2016) |
| HEI193 | Schwannoma derived from peripheral nervous system/vestibular nerve of a NF2 patient, transformed cell line | *NF2:* c.15751G>A (Splice acceptor mutation/splicing defect) | | Round | (Hung et al., 2002) |
| sNF96.2 | MPNST | *NF1:* c.3683delC (p.Asn1229Metfs*11) | *NF1* LOH | Spindle | (Li et al., 2004) |
| sNF02.2 | MPNST | NF1 | | Spindle | (Li et al., 2004) |
| STS26T | MPNST derived from the metastatic site of bone at left scapula | No NF1/*NF1* wild type | Homozygous deletion of *TP53* | Spindle | (Dahlberg et al., 1993) |
| T265 | MPNST | NF1 | | Spindle | (Badache and De Vries, 1998) |
| ST8814 | MPNST | *NF1:* c.910C>T (p.Arg304Ter) | *NF1* LOH | Spindle | (Glover et al., 1991) |
| S462 | MPNST derived from thigh | *NF1:* c.6792C>A (p.Tyr2285Ter) *TP53:* c.329G>C (p.Arg110Pro) | *NF1* LOH, *TP53* LOH | Spindle | (Frahm et al., 2004) |
| RHT163 | MPNST | NF1 | N/A | Round | (Gampala et al., 2024) |
| RHT172 | MPNST | NF1 | N/A | Round | (Gampala et al., 2024) |
| RHT92 | MPNST PDX cell line/xenoline | NF1 | N/A | Spindle | (Gampala et al., 2021) |
| JH-2-079c | MPNST | NF1 | *NF1-/-* | Epithelial/variable (round) | (Pollard et al., 2020) |
| JH-2-002 | MPNST | NF1 | *NF1-/-* | Spindle | (Pollard et al., 2020) |
| JH-2-103 | MPNST | NF1 | *NF1-/-* | Spindle | (Pollard et al., 2020) |

**Table S4. The information on the human cell Lines.**
